# Integrated coding-noncoding genome annotation expands single-cell transcriptomic discovery and identifies clinically relevant noncoding RNAs in multiple myeloma

**DOI:** 10.64898/2026.08.09.743753

**Authors:** Marina E. Michaud, Denis J. Ohlstrom, Mojtaba Bakhtiari, Edward Henderson, Sarthak Satpathy, Katherine E. Ferguson, William C. Pilcher, Edgar Gonzalez-Kozlova, Dimitra Karagkouni, Shannon M. Matulis, Chaitanya R. Acharya, MMRF Immune Atlas Consortium, David Avigan, Ravi Vij, Samir Parekh, Hearn Jay Cho, Ioannis S. Vlachos, Li Ding, Shaji Kumar, Sacha Gnjatic, Ajay Nooka, George Mulligan, Sagar Lonial, Lawrence H. Boise, Manoj Bhasin

## Abstract

Although the human genome encodes a vast repertoire of noncoding RNAs that regulate gene expression, the noncoding genome remains underexplored due to technical challenges. Specifically, during transcriptomic sequencing data alignment, the overlap between noncoding and coding loci can create ambiguous read alignments that are subsequently discarded from downstream analysis. For this reason, most of the noncoding genome is excluded from standard genomic annotations used for sequencing alignment. To address this challenge and enable concurrent profiling of the coding and noncoding transcriptome, we systematically integrated standard coding (GENCODE) and noncoding (LncBook) genome annotations, preserving coding gene annotations and removing overlapping noncoding regions. The resulting integrated genome annotation expanded the number of annotated noncoding genes from 40,785 to 138,296 while preserving all coding genes and reducing ambiguous read assignment. To evaluate the utility of our integrated genome annotation for uncovering novel, biologically relevant noncoding RNAs (ncRNAs), we realigned CD138-positive bulk RNA-seq (N = 942) and CD138-negative single-cell RNA-seq (N = 478) data from the MMRF CoMMpass study, generating a comprehensive coding-noncoding atlas of the myeloma bone marrow microenvironment with noncoding genes representing 51% of highly variable genes and displaying significant cell type specificity. Tumor expression profiling based on this integrated profiling identified 15 clusters, including two enriched for amp(1q21) or t(4;14) and associated with shorter progression-free survival (PFS). Differential expression and systematic filtering yielded 19 candidate high-risk ncRNAs, including previously uncharacterized *ENSG00000310209*, which was associated with poor PFS (HR = 1.141, *P* = 0.0025), increased IRF4 activity, Wnt pathway activation, CCL5 signaling, and the accumulation of anergic-like CD8+ T cells. These findings establish integrated coding-noncoding analysis as a strategic approach for discovering functional ncRNAs from transcriptomic sequencing data.

## Introduction

While the protein-coding genome has long dominated genomic investigations, it represents only a small fraction of the human transcriptome. The majority of the genome is instead defined by a vast noncoding landscape comprising tens of thousands of RNAs whose functions remain largely unknown; however, mounting evidence indicates that many play crucial roles in epigenetic, transcriptomic, and proteomic regulation of both physiological and pathological processes.^1^ Specifically, noncoding RNAs (ncRNAs) have been implicated in malignancy-driving processes such as cell-state control,^2,3^ lineage plasticity,^4,5^ oncogenic signaling,^6,7^ immune evasion,^8,9^ and treatment response,^10–12^ suggesting that the noncoding transcriptome is not merely an ancillary genomic component, but a substantial and largely untapped facet of cancer biology. Its systematic investigation, therefore, offers a major opportunity to uncover novel drivers of tumorigenesis and disease progression, potentially refining prognostication and therapies.

Despite this potential, large-scale investigation of the noncoding genome remains hampered by technical challenges. Standard reference annotations, such as GENCODE and Ensembl,^13,14^ prioritize well-supported genes and thus capture only a limited subset of the noncoding transcriptome. Conversely, dedicated noncoding reference annotations, such as LncBook,^15^ contain orders of magnitude more noncoding gene annotations by incorporating predicted noncoding transcripts from various databases. However, the isolation of these annotations leads to parallel analyses of coding and noncoding genes, limiting the biological interpretation of noncoding genes and their relationships to protein-coding pathways, and underscoring the need for integrated coding-noncoding analyses. Furthermore, because many noncoding transcripts genomically overlap with coding loci, the simple concatenation of independent reference annotations creates significant ambiguity in read assignment, compromising quantification and downstream analyses.

Consequently, a wealth of existing sequencing data, including large-scale cancer genomic studies, cannot be robustly mined for noncoding genes using standard analytical approaches. Here, we address this challenge by generating an integrated coding-noncoding genome annotation designed for comprehensive transcriptomic analysis from standard sequencing data. Our approach systematically resolves conflicting genomic coordinates between coding and noncoding loci by prioritizing mRNA annotations while trimming overlapping ncRNA regions. This reduces ambiguous read mapping without sacrificing noncoding coverage, enabling alignment of existing and new bulk and single-cell RNA-sequencing datasets to provide simultaneous insight into coding and noncoding gene expression. By doing so, this approach opens the large body of existing and future datasets to interrogation of the previously obscured noncoding landscape that governs cell states and hallmark pathways.

As a proof of concept, we applied this approach to investigate multiple myeloma (MM), a plasma cell malignancy that remains marked by high relapse rates, treatment resistance, and poor survival despite clinical advances over the past two decades.^16^ While cytogenetic and clinical features help identify patients with high-risk disease, the prediction of rapid disease progression and poor treatment response still poses a significant challenge.^17^ By applying integrated coding-noncoding analysis to the Multiple Myeloma Research Foundation (MMRF) CoMMpass study RNA sequencing datasets, we uncovered ncRNAs linking tumor phenotype to disease progression, immune dysfunction, and therapeutic response. Specifically, using our integrated genome annotation, we profiled coding and noncoding transcription across the myeloma bone marrow microenvironment (BMME), identifying 19 candidate ncRNAs of interest. Collectively, our results demonstrate that integrated analysis of coding and noncoding genes reveals biologically relevant features hidden by standard approaches, expanding the interpretable cancer transcriptome and creating new opportunities for biomarker discovery, mechanistic investigation, and therapeutic development.

## Results

### Systematic integration of standard and noncoding genome annotations enables comprehensive profiling of the coding and noncoding genome

To enable concurrent analysis of coding and noncoding transcription from RNA-sequencing data, we developed a systematic workflow to merge standard and noncoding genome annotations into a single, integrated reference. Direct concatenation of these annotations is problematic due to the overlap of sense and antisense ncRNA transcripts with coding exons (Figure 1a). Such conflicts create ambiguous alignments, in which reads mapping to shared genomic intervals cannot be confidently assigned and are subsequently discarded, limiting transcript capture and complicating joint analysis of coding and noncoding genes.

**Figure 1.**
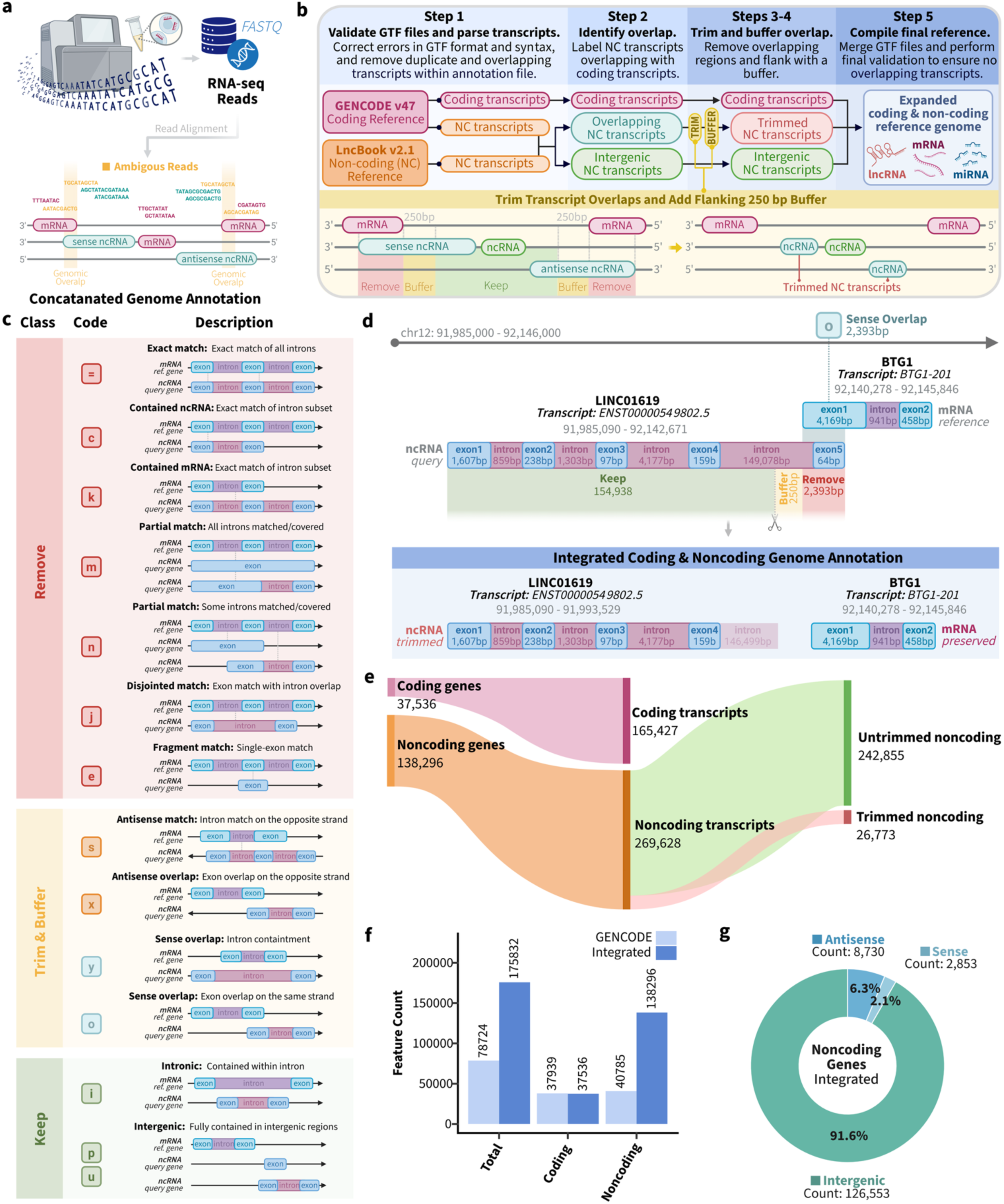
Integration of coding and noncoding genome annotations for concurrent transcriptomic profiling. **(a)** Overview of how overlaps in coding and noncoding gene annotations result in ambiguous reads during read alignment of raw sequencing data. **(b)** Schematic of the annotation integration pipeline used to merge GENCODE v47 and LncBook v2.1 into a unified coding-noncoding reference. Overlapping noncoding transcript models were resolved by trimming shared regions and removing an additional 250-bp flanking buffer to reduce downstream alignment ambiguity. **(c)** Classification of transcript overlap patterns annotated during Step 2 of the integration pipeline. Noncoding transcripts were annotated according to overlap with coding genes and either completely removed, trimmed and buffered in Step 3, or preserved. **(d)** Representative genomic locus illustrating annotation integration of the noncoding transcript LINC01619, which displays sense overlap with the coding transcript BTG1. In the integrated reference, the overlapping portion of the noncoding transcript and the 250-bp flanking buffer were removed, preserving the coding annotation while retaining the non-overlapping segment of the ncRNA transcript. **(e)** Sankey diagram displaying the number of coding and noncoding genes in the final integrated reference genome annotation. Genes are composed of preserved coding, untrimmed noncoding, and trimmed noncoding transcripts. **(f)** Comparison of coding and noncoding gene content between the integrated reference and GENCODE v47. **(g)** Composition of noncoding genes in the integrated reference, stratified as sense, antisense, and intergenic.

To overcome this problem, we established a pipeline with five steps to resolve duplicate and overlapping gene annotations and compile a final integrated coding-noncoding reference annotation (Figure 1b). We applied this approach to GENCODE v47 and LncBook v2.1, a comprehensive noncoding annotation comprising 526,318 ncRNA transcripts aggregated from ten noncoding reference databases.^15^ The pipeline first parses and cleans the input GTF files, separating coding and noncoding transcripts, removing duplicate entries, and consolidating fragmented transcripts into unified loci. The transcripts of the clean GTF files are then directly compared to assess the overlap of noncoding and coding transcripts, assigning class codes to each transcript (Figure 1c). During this process, all coding transcripts are preserved in full, as are noncoding transcripts that do not overlap with coding loci (i.e., classes “i”, “p”, and “u”). By contrast, noncoding transcripts that display either sense or antisense overlap with coding transcripts (i.e., classes “s”, “x”, “y”, and “o”) are systematically trimmed to remove the shared region. To further reduce ambiguity at coding-noncoding boundaries, we additionally remove a 250 base-pair (bp) flanking buffer from the affected end(s) of the noncoding transcript (Figure 1b). For example, a transcript of the long noncoding RNA (lncRNA) gene *LINC01619* displays sense overlap with a transcript of the coding gene *BTG1* on chromosome 12 (Figure 1d). The pipeline identifies and annotates this sense overlap and removes the 2,393-bp overlapping region, along with an additional 250-bp buffer from the noncoding transcript, which, in this case, creates a terminal intronic sequence that is not present in the final integrated annotation. Notably, this trimming process affected only 10% of ncRNA transcripts present in the final integrated reference (Figure 1e). This conservative strategy preserves the integrity of the coding annotation while retaining as much noncoding information as possible for downstream quantification.

Collectively, the final integrated reference retained the original coding genes while markedly expanding the accessible noncoding landscape, yielding 175,832 unique annotated genes, of which 78.6% were noncoding (Figure 1e), including predominantly intergenic noncoding genes (Figure 1f). This represented a more than twofold increase in retained features relative to GENCODE alone (78,724 total genes) and markedly expanded detection of sense, antisense, and intergenic ncRNAs from 40,785 to 138,296 genes, providing a broader and more comprehensive representation of the expressed transcriptome. These results indicate that systematic integration can substantially broaden transcriptome coverage without disrupting the core coding annotation.

### Benchmarking the performance of the integrated genome annotation against standard and concatenated annotations

We next evaluated the performance of the integrated reference by separately realigning a single-cell RNA-seq benchmarking dataset of myeloma BM samples (N = 48) to GENCODE v47, a concatenation of GENCODE v47 and LncBook v.2.1, and our integrated reference annotation to test whether our approach: (1) reduced ambiguous read alignment, (2) improved noncoding gene capture without sacrificing coding gene capture, and (3) resulted in high-quality ncRNAs guiding clustering and cell type annotation (Figure 2a). After filtering for high-quality features with more than 10 read counts and detection in more than 20% of samples, alignment to the integrated reference expanded the number of captured noncoding features from 28,088 to 70,571 (Figure S1A). Importantly, noncoding gene capture did not occur at the expense of coding genes, with 26,218 coding features captured using the integrated reference genome, compared with 25,724 using GENCODE v47 (Figure S1A). Cumulatively, these findings indicate that the integrated annotation expands the explorable transcriptome rather than redistributing reads away from well-characterized mRNAs.

**Figure 2.**
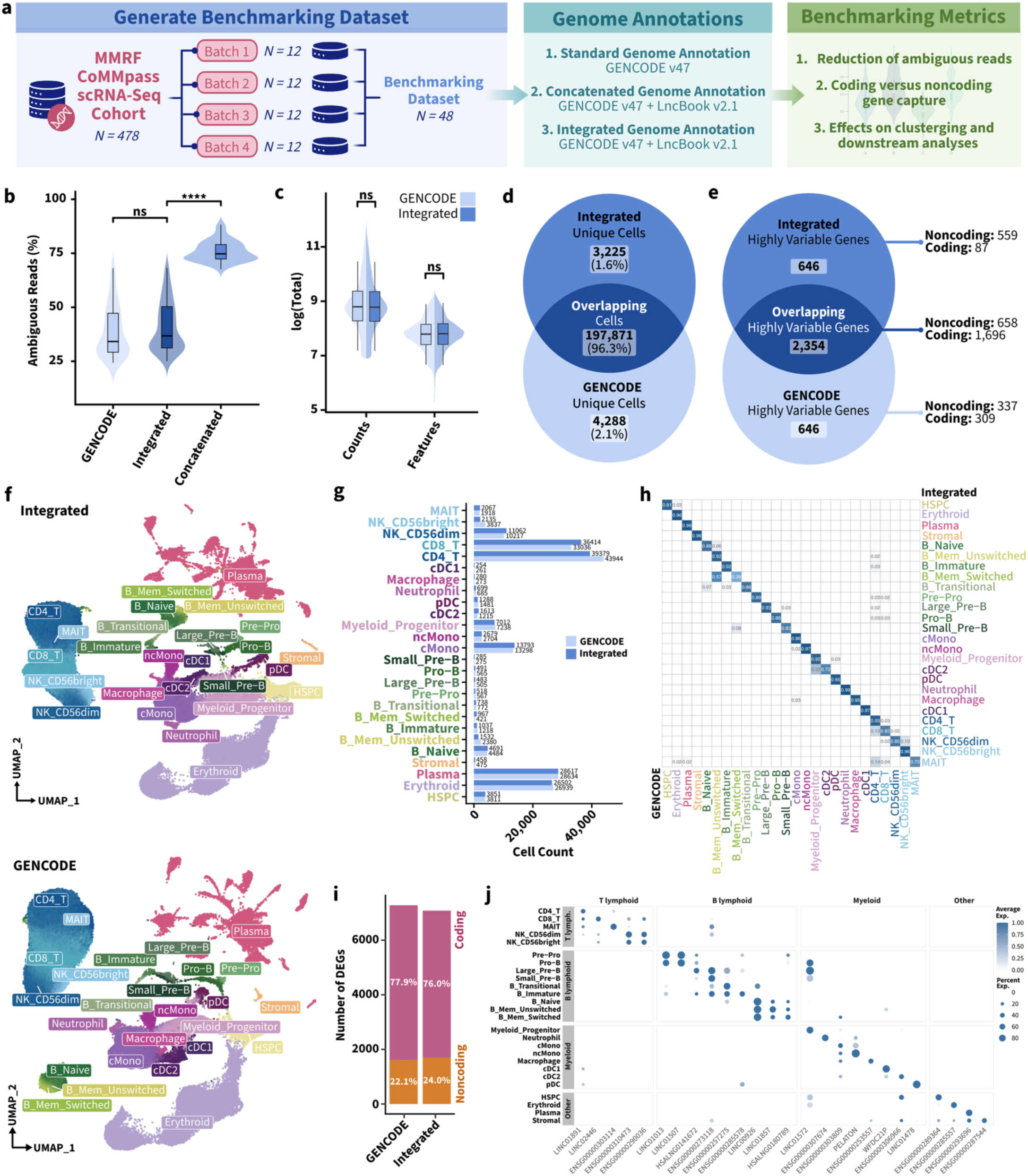
The integrated coding and noncoding genome annotation reduces ambiguous reads while improving the capture of significant ncRNAs. (a) Overview of how the scRNA-seq benchmarking dataset was generated to evaluate our integrated genome annotation against the standard, concatenated, and integrated genome annotations. (b) Violin plots displaying the percentage of ambiguous reads following alignment with GENCODE v47, the integrated reference, and the concatenated GENCODE v47 and LncBook v2.1 reference. Comparisons were performed using paired Wilcoxon signed-rank tests with Holm correction (\*\*\*\**P* < 0.0001). (c) Split violin plots showing the number of counts and detected features per cell after quality control (QC) filtering. Alignment to the integrated reference increased the number of detected features due to expanded noncoding coverage, without significantly reducing average counts per cell. Comparisons were performed using paired Wilcoxon signed-rank tests with Holm correction. (d) Venn diagram illustrates the overlap of barcoded cells passing quality-control thresholds recovered when aligning to GENCODE v47 versus the integrated reference. While most cells were present in the final, filtered count matrices following both alignments, some cells were exclusively recovered after alignment to the integrated genome annotation or GENCODE v47 annotation. (e) Overlap among the top 3,000 highly variable genes (HVGs) identified from each alignment, showing increased capture of noncoding HVGs with the integrated reference. (f) UMAP projection of major BMME cell populations identified using identical clustering parameters in datasets aligned to GENCODE or the integrated reference. (g) Comparison of cell-type abundance between the two aligned datasets. (h) Confusion matrix showing concordance of cell-type assignments among barcoded cells present in both the GENCODE-and integrated-aligned datasets. (i) Proportion of coding and noncoding DEGs across cell populations (absolute log_2_fold-change > 3.0, detection > 10%, *P-adj.* < 0.05). (j) Cell-type-specific expression of representative noncoding genes across the multiple myeloma bone marrow microenvironment. Dot size indicates the percentage of cells expressing each gene, and color indicates average scaled expression.

Because a central goal of our approach was to avoid read ambiguity, we next compared our integrated genome annotation against a concatenated reference generated by merging GENCODE v47 with LncBook v2.1 without resolving overlapping loci. The proportion of ambiguous reads was substantially reduced from a median of 74.8% ambiguous reads per-sample using the concatenated reference to 34.2% using our integrated reference, underscoring the importance of systematically resolving overlaps rather than simply combining GTF files (*P* = 2.22×10^-16^, Figure 2b). As expected, the average number of unique features captured per cell increased marginally, from a median of 2,417 to 2,454, due to the expanded noncoding annotation. A theoretical limitation of this approach is that average count detection may decrease due to ncRNAs typically being expressed at lower levels than coding transcripts;^18–20^ however, we found that the median number of counts per cell (N = 6,509) remained highly similar to GENCODE (N = 6,604; Figure 2c).

At the cellular level, the two alignments were highly concordant. Overall, 96% of cells passing quality control were shared between the datasets aligned using the GENCODE versus integrated references, with only 2% of cells being unique to either annotation (Figure 2d). These mutually exclusive cells did not differ in standard quality control metrics (Figure S1B), suggesting that low-quality noise was not the source of annotation-specific cell capture. Instead, the identities of these unique cell sets suggest that transcriptome representation may influence the recovery of specific cell states with some cellular subtypes being exclusively represented (Figure S1C).

We next asked whether the newly captured noncoding features altered downstream analysis. In the dataset aligned to the integrated annotation, noncoding genes were well represented among highly variable genes rather than being confined to low-information background features (Figure 2e), indicating that they contribute to the transcriptional heterogeneity that drives single-cell clustering. Consistent with this, clustering of the GENCODE-and integrated-aligned datasets using identical parameters preserved the overall architecture of the MM-BMME and recovered the same major immune and stromal populations (Figure 2f, g). Direct comparison of cellular annotations between the two datasets showed high concordance (>90%) across most populations (Figure 2h). However, several compartments, namely myeloid progenitors, memory B, CD4 T, CD8 T, NK, and MAIT cells, showed greater divergence between annotations. While we hypothesize that the expanded number of features introduced by alignment to the integrated reference refines clustering and cell-type assignment in selected compartments, the absence of cell-specific ground-truth labels makes it difficult to determine which alignment is most accurate for these subsets.

To further probe whether ncRNAs contributed to cell-type discrimination, we examined the significantly differentially expressed genes (DEGs) across clusters. Noncoding genes accounted for approximately 24% of total DEGs (Figure 2i) and exhibited high cell-type specificity (Figure 2j), indicating that they are not only detectable but informative for defining cellular identity and state. Collectively, these findings show that systematic integration of coding and noncoding annotations enables robust joint transcriptomic analysis, expands detectable transcriptional space without compromising quality, and reveals a new layer of biologically informative noncoding transcription in the MM-BMME.

### Coding-noncoding single-cell atlas of the myeloma bone marrow microenvironment

Having established that integrated coding-noncoding alignment expands the capture of biologically informative noncoding transcripts, we next applied this reference to construct a single-cell atlas of the coding-noncoding transcriptome in the multiple myeloma bone marrow microenvironment. Because the immune microenvironment (IME) is a major determinant of disease progression and therapeutic response in MM,^21–23^ we sought to test whether expanded ncRNA capture would improve the resolution of cellular states shaping patient outcomes.

To this end, we realigned CD138-negative bone marrow aspirate samples (N = 478) from 337 patients enrolled in the MMRF CoMMpass study to the integrated genome annotation (Figure 3a). The resulting atlas comprised 2,092,670 high-quality cells spanning lymphoid, myeloid, erythroid, stromal, and plasma cell lineages (Figure 3b, Figure S2). The presence of plasma cells within the CD138-negative fraction was consistent with incomplete depletion in a subset of samples and provided an internal reference for malignant plasma cell transcriptional states. Following quality-control filtering, 96,789 features with more than 10 read counts and detection in more than 20% of samples were retained in the final count matrix, of which 77% corresponded to noncoding genes (Figure 3c). Although these ncRNA features accounted for only 8% of total captured counts and ∼51% of highly variable genes (Figure 3c), their broad representation is consistent with the large breadth of the noncoding transcriptome and the generally lower abundance of ncRNAs relative to coding transcripts.^18–20^

**Figure 3.**
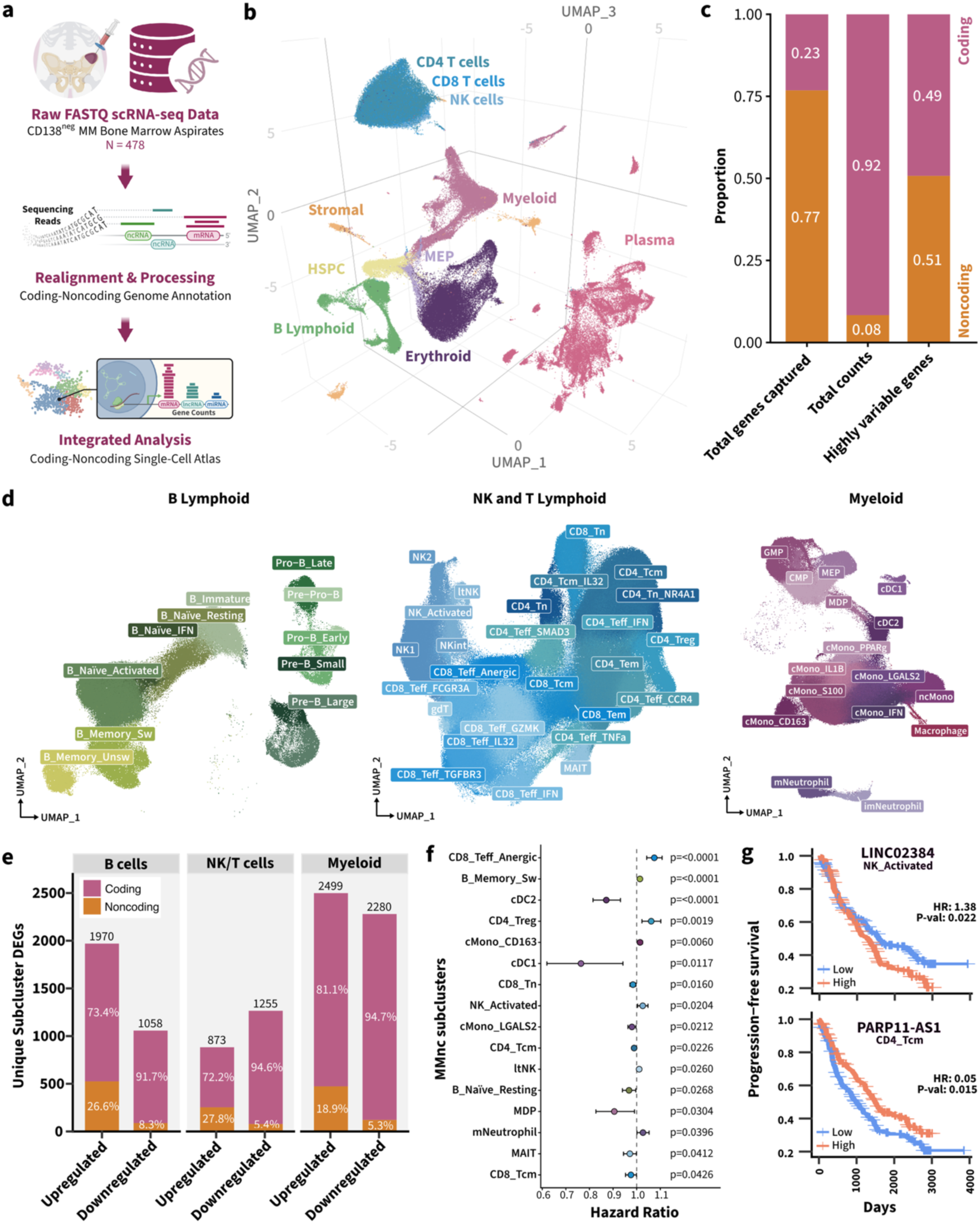
Coding-noncoding single-cell atlas of the myeloma bone marrow microenvironment. (a) Overview of the analytical workflow used to generate the coding-noncoding single-cell atlas from CD138-negative bone marrow aspirate samples. (b) Three-dimensional UMAP projection of the resulting atlas showing the major cellular compartments of the multiple myeloma bone marrow microenvironment. (c) Stacked bar plots display the proportion of coding and noncoding genes among total genes captured, total transcript counts, and highly variable genes in the integrated atlas passing quality control with more than 10 total counts, detected in more than 10 cells, and expressed in more than 20% of samples. (d) UMAP projections of subclustered B lymphoid, NK/T lymphoid, and myeloid compartments, yielding 52 distinct immune subpopulations across the atlas. (e) Proportion of coding and noncoding unique DEGs (log_2_fold-change <-1.0 or > 2.0, detection > 10%, *P-adj.* < 0.05) among upregulated and downregulated subcluster markers within the B cell, NK/T cell, and myeloid compartments. Numbers above bars indicate the total number of unique DEGs in each category. (f) Forest plot showing associations between immune subcluster abundance (as a proportion of each subcluster within myeloid, B lymphoid, NK cell, CD4+ cell, and CD8+ T cell compartments) and progression-free survival. Points represent hazard ratios per unit increase in subcluster abundance from Cox proportional hazards models, with error bars indicating 95% confidence intervals and significance determined using a two-sided Wald test. (g) Kaplan-Meier curves visualizing representative associations between subcluster-specific ncRNA expression and PFS. High-and low-expression groups were defined using an optimal cut point for visualization only. Hazard ratios and P values were derived from Cox proportional-hazards models using per-patient pseudobulk ncRNA expression as a continuous predictor. High abundance of *LINC02384* in activated NK cells was associated with worse outcomes, whereas high abundance of *PARP11-AS1* in CD4+ central memory T cells was associated with better outcomes.

We next performed compartment-level subclustering to resolve finer immune cell states within the B lymphoid, NK and T lymphoid, and myeloid compartments, identifying a total of 52 subpopulations (Figure 3d, Figure S3 and S4). These included canonical developmental and immune states as well as transcriptionally distinct interferon-responsive populations within multiple compartments, including B_Naïve_IFN, CD4_Teff_IFN, CD8_Teff_IFN, and cMono_IFN. The recurrence of these interferon-stimulated subpopulations across lineages aligns with previous findings that interferon signaling is a prominent feature shaping the MM-BMME and patient outcomes.^24–26^ We then asked whether ncRNAs contributed meaningfully to the transcriptional identity of these immune states. Among unique subcluster markers, noncoding genes accounted for up to 27.8% of upregulated and 8.3% of downregulated genes, demonstrating that ncRNAs are prominent components of the transcriptional profiles defining immune cell identity and state (Figure 3e).

To determine whether these immune states were clinically relevant, we next evaluated associations between subcluster abundance and patient outcomes. This analysis identified 16 immune subpopulations significantly associated with patient outcomes (*P* < 0.05, Figure 3f, Figure S5). Within the B cell compartment, poor outcomes were associated with a lower abundance of resting naïve B cells (HR = 0.968, *P* = 0.027) and an increased abundance of switched memory B cells (HR = 1.014, *P* = 0. 1.57×10^-5^), consistent with a more antigen-experienced immune repertoire,^27,28^ potentially related to immune aging and myelomagenesis. Within the T cell compartment, reduced MAIT cell abundance was associated with poor outcomes (HR = 0.971, *P* = 0.041), consistent with prior reports linking MAIT cell loss to aging, immune dysfunction, and MM progression.^29,30^ Within the myeloid compartment, decreased abundance of mature (cDC1: HR = 0.764, *P* = 0.012; cDC2: HR = 0.871, *P* = 5.1×10^-5^) and immature (HR = 0.905, *P* = 0.030) classical dendritic cell populations was also associated with poor outcomes, suggesting that depletion of antigen-presenting cells may contribute to impaired anti-tumor immunity. Collectively, these findings recapitulate established immune correlates of poor outcome in MM and provide a clinically relevant foundation for identifying noncoding RNAs associated with these adverse immune states.

We then examined the association of representative ncRNA markers enriched in these outcome-associated subclusters with progression-free survival (PFS). Among the top ncRNA subcluster DEGs associated with outcomes, increased expression of *LINC02384* in activated NK cells (HR = 1.38, *P* = 0.022) and decreased expression of *PARP11-AS1* in CD4^+^ central memory T cells (HR = 0.05, *P* = 0.015) were both linked to poor outcomes (Figure 3g). Interestingly, *LINC02384* has been previously associated with altered immune responses, including positive co-expression with IFN-γ expression.^31,32^ Consistent with this association, activated NK cells highly expressing *LINC02384* in our dataset exhibited significantly increased IFN-γ expression (*P* = 0.024, Figure S6). Additionally, *PARP11* expression has been shown to promote T_reg_ accumulation and an immunosuppressive TME, suggesting that its antisense ncRNA counterpart, *PARP11-AS1,* may putatively counteract these effects.^33,34^ Taken together, this data establishes a coding-noncoding single-cell atlas of the MM bone marrow microenvironment and demonstrates that ncRNAs are integral features of clinically relevant immune states, providing a foundation for the subsequent identification of noncoding genes associated with high-risk myeloma phenotypes.

### Integrated bulk and single-cell profiling identifies prognostic tumor-intrinsic ncRNAs associated with high-risk myeloma phenotypes

We first sought to identify tumor-associated ncRNAs linked to high-risk disease. Building on the known associations among ncRNAs, tumor progression, and outcomes in MM,^35^ we hypothesized that high-risk disease is, in part, shaped by ncRNA-driven myeloma cell phenotypes. We therefore next asked whether high-risk myeloma samples exhibit distinct noncoding profiles associated with specific phenotypes and poor outcomes.

To address this, we realigned bulk RNA-seq data from the CD138-positive fractions of the MMRF CoMMpass bone marrow samples to the integrated genome annotation (N = 942; Figure 4a). Although the CD138-negative fractions used to generate the single-cell atlas contained residual plasma cells in a subset of samples, likely owing to incomplete depletion in patients with higher tumor burden (ρ = 0.8, *P* = 1.783×10^-101^, Figure S7), the CD138-positive bulk cohort provided broader patient coverage and more robust representation of tumor-intrinsic transcriptional programs. We therefore used the CD138-positive bulk RNA-seq data for primary discovery of myeloma-associated ncRNAs, while leveraging the CD138-negative single-cell atlas to confirm plasma-cell-specific expression and to define associated immune microenvironmental features.

**Figure 4.**
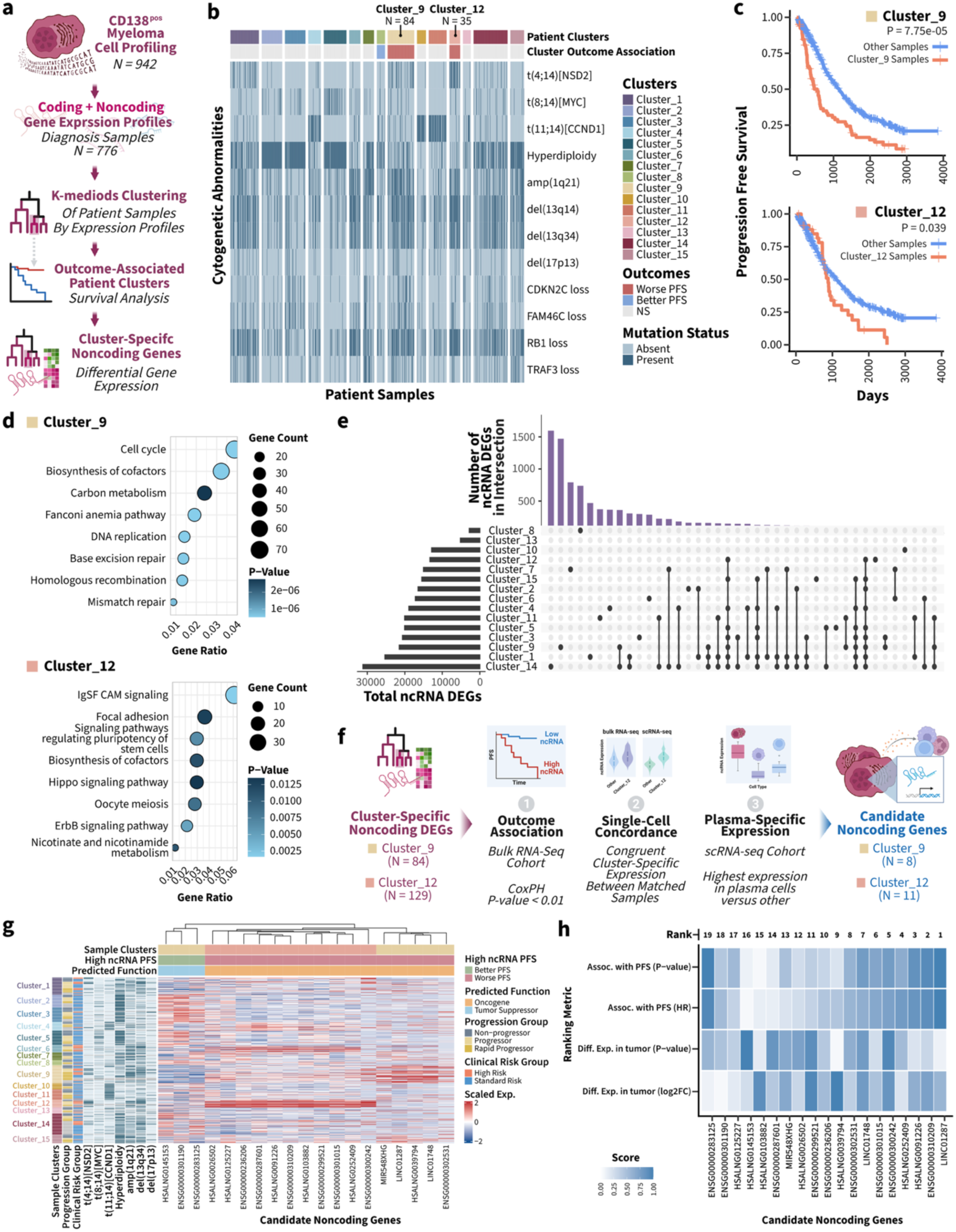
Integrated bulk and single-cell profiling identifies prognostic tumor-intrinsic ncRNAs associated with high-risk myeloma phenotypes. **(a)** Analytical workflow for discovery of cluster-specific noncoding genes from CD138-positive myeloma bulk RNA-seq profiles. Bulk RNA-seq data from 942 CD138-positive myeloma-enriched fractions, representing 776 diagnosis samples, were realigned to the integrated coding-noncoding genome annotation, clustered by transcriptomic similarity, and tested for associations with clinical outcome to identify cluster-specific noncoding genes. **(b)** Heatmap showing the fifteen patient clusters generated by consensus clustering of patient coding-noncoding transcriptomes with the distribution of cytogenetic abnormalities across samples. Cluster 9 and Cluster 12 were enriched for high-risk cytogenetic features, including amp(1q21) and t(4;14), respectively. **(c)** Kaplan-Meier plots showing worse PFS for patients in Cluster 9 and Cluster 12 compared with all other samples using multivariable Cox proportional hazards models with a two-sided Wald test. **(d)** KEGG pathway enrichment analysis of genes upregulated in Cluster 9 and Cluster 12 relative to other clusters, with pathway significance reported using a hypergeometric over-representation test. Dots are sized by the number of differentially expressed genes present in the respective pathway and colored by the raw *P*-value for pathway enrichment. **(e)** UpSet plot summarizing the number and overlap of ncRNA DEGs across patient clusters. Horizontal bar plot indicates the total number of ncRNA DEGs per cluster, and connected dots indicate shared or cluster-specific intersections. **(f)** Multistep strategy used to refine candidate ncRNAs from high-risk patient clusters. Cluster-specific ncRNAs were filtered by association with PFS in the bulk RNA-seq cohort, concordant cluster-specific expression in matched bulk and single-cell samples, and preferential expression in plasma cells in the single-cell atlas. **(g)** Heatmap of candidate ncRNAs derived from Cluster 9 and Cluster 12, showing their expression across patient samples grouped by cluster on the y-axis with associated progression group, clinical risk, and cytogenetic abnormalities. Hierarchical clustering of candidate ncRNAs shows similar expression profiles for downregulated ncRNAs associated with better PFS (i.e., predicted tumor suppressors) and upregulated ncRNAs associated with worse PFS (i.e., predicted oncogenes). **(h)** Ranking of candidate ncRNAs based on differential expression, association with PFS, and relationships to altered abundance of outcome-associated immune subpopulations. Significant differential expression was determined using the DESeq2 Wald test with Benjamini-Hochberg correction, while significant outcome association was determined using multivariable Cox proportional hazards models with a Wald test. The heatmap shows scaled ranks across the indicated metrics, with lower composite rank indicating higher overall priority, for the top 10 candidate ncRNA-immune subpopulation interactions.

Although cytogenetic features remain clinical hallmarks for risk stratification, their inherent heterogeneity among patients complicates the analysis of independent events. Recent evidence suggests that expression profile-and outcome-based patient stratification enables more robust identification of prognostic features from transcriptomic data.^23,36^ Therefore, to identify high-risk patients, we performed consensus clustering of the bulk tumor transcriptomes, revealing 15 distinct patient clusters (Figure 4a, b). Although these clusters were defined solely by coding and noncoding gene expression, they showed discernible trends in cytogenetic composition, similar to expression profile-derived patient clusters reported by Skerget *et al.* (Figure 4c, Figure S8). Most notably, Cluster 9 and Cluster 12 were enriched for amp(1q21) and t(4;14), respectively, and both were associated with worse progression-free survival (Cluster 9: HR = 1.73, *P* = 7.75e-05; Cluster 12: HR = 1.58, *P* = 0.039).

To better define the biology of these outcome-associated clusters, we next examined the pathways enriched in each cluster (Figure 4d). Cluster 9 was characterized by the upregulation of cell cycle (e.g., *CCNB1*, *CDK1, E2F2*) and DNA replication (e.g., *MCM2*, *POLD1*, *FEN1*) genes (Table S3), consistent with a highly proliferative, more aggressive phenotype, as reported previously by Skerget *et al.* (Figure S8).^36^ By contrast, Cluster 12 was marked by increased expression of immunoglobulin superfamily cell-adhesion molecules (Table S4), including *PVR* and *JAM3*, implicating pathways linked to immune dysregulation and disease progression.^37,38^ Together, these data indicated that poor-outcome transcriptomic clusters capture biologically distinct high-risk myeloma phenotypes.

Next, we sought to determine whether these high-risk clusters exhibit unique noncoding gene expression patterns driving their aggressive phenotype. Differential expression analysis identified cluster-specific ncRNAs across the 15 patient groups, with several clusters, including Clusters 9 and 12, showing substantial proportions of unique noncoding DEGs (Figure 4e). To isolate the most informative candidates, we removed ncRNAs that were recurrently upregulated or downregulated across multiple clusters and selected those with high baseline expression, yielding 1,517 unique cluster-specific ncRNAs across the 15 clusters. We then applied a multistep filtering workflow to select for ncRNAs most likely to be biologically and clinically relevant (Figure 4f).

Because Clusters 9 and 12 were most strongly associated with poor progression-free survival, we focused subsequent analyses on ncRNAs specific to these two groups. This yielded 213 cluster-specific ncRNAs, which we then tested for association with progression-free survival independent of cluster membership. Seventy-two ncRNAs remained significantly associated with outcome (*P* < 0.01). To further refine this list, we leveraged the single-cell cohort and examined matched samples shared between the bulk and single-cell datasets (N = 254), asking whether candidate ncRNAs retained concordant cluster specificity and showed preferential expression in plasma cells within the single-cell atlas (Figure S8). This filtering strategy yielded 19 top candidate ncRNAs that satisfied all quality-control criteria (Figure 4g).

Lastly, to prioritize the candidate ncRNAs, we ranked them based on their differential expression and association with outcomes (Figure 4h). Interestingly, the first-ranked ncRNA candidate, *LINC01287,* has been implicated in the progression of breast, lung, and liver cancers,^39–41^ but has not been previously characterized in myeloma. Consistent with these reports, high tumor-intrinsic *LINC01287* expression was associated with increased expression of previously described LINC01287-linked mRNA targets, including *IGF1R*, *WNT5A*, and *TCF12*, in our cohort (Figure S9). Thus, our workflow and prioritization strategy recovered a cancer-associated ncRNA with conserved target relationships, supporting its ability to identify biologically relevant noncoding candidates in high-risk myeloma.

In addition to known cancer-associated ncRNAs, the candidate list included several uncharacterized noncoding genes. We therefore focused subsequent analyses on *ENSG00000310209*, hereafter referred to as nc209, the second-ranked candidate with no established role in myeloma or cancer biology, allowing us to test whether our integrated coding-noncoding profiling approach can move beyond rediscovery of known cancer ncRNAs to nominate previously unrecognized regulators of high-risk disease.

### nc209 is a novel ncRNA in high-risk myeloma linked to Wnt pathway activation, IRF4 activity, and CCL5 signaling

Because nc209 is uncharacterized, we sought to infer its potential function through co-expression and inter-and intracellular signaling analyses, validating our findings *in vitro*.

We first asked whether nc209 showed cytogenetic specificity consistent with the high-risk transcriptomic cluster from which it was nominated. Comparing three MM cell lines, nc209 expression was higher in those bearing t(4;14) translocation than those without (Figure 5a), mirroring its enrichment in Cluster 12, which was characterized by increased t(4;14) prevalence and inferior PFS. These findings support nc209 as a candidate ncRNA specific to a high-risk myeloma cell state.

**Figure 5.**
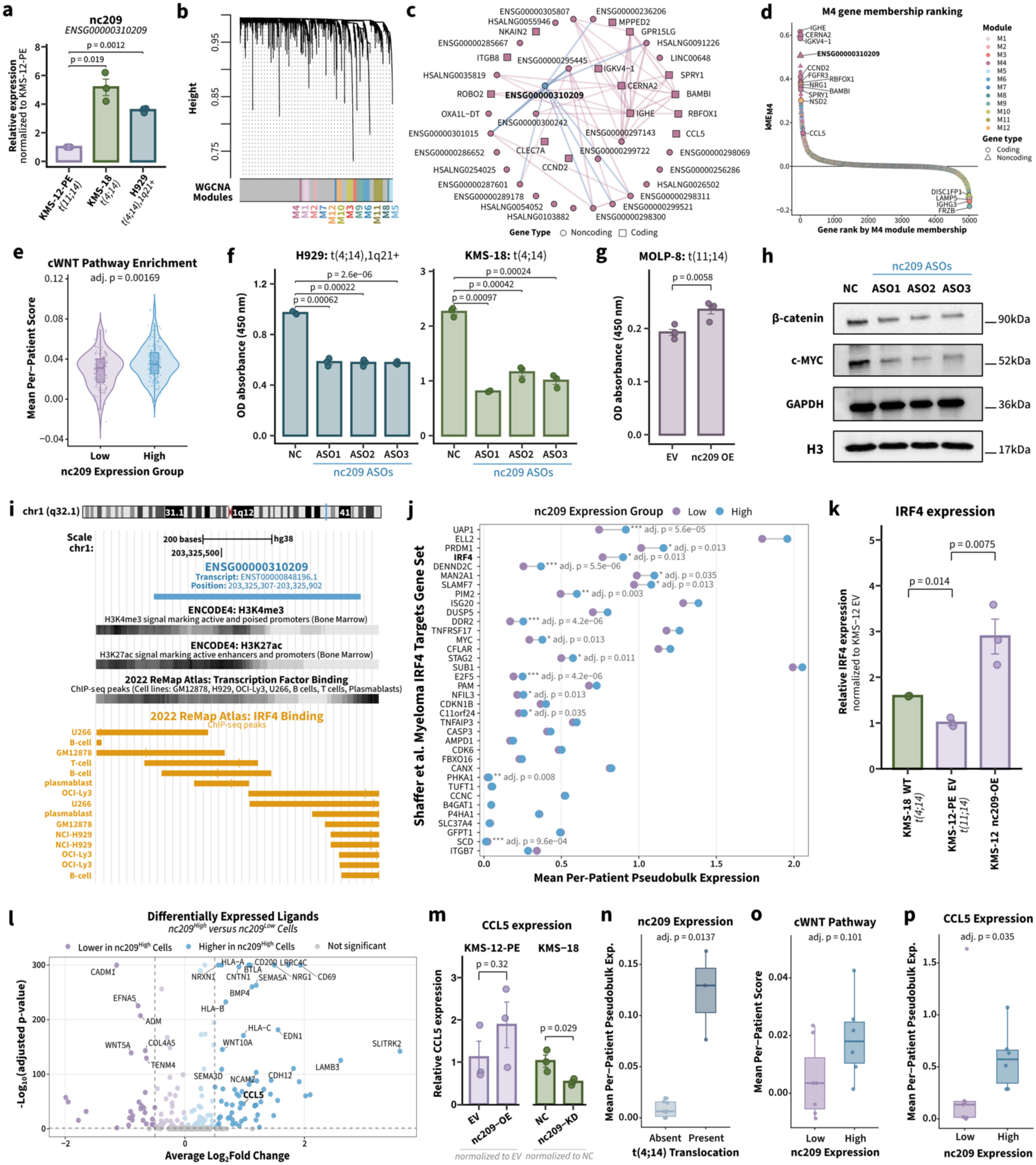
nc209 is a t(4;14)-associated ncRNA linked to Wnt pathway activation, IRF4 activity, CCL5 expression, and myeloma cell fitness. **(a)** KMS-18 and H929 cell lines harboring t(4;14) showed higher relative expression of *ENSG00000310209* (nc209) measured by RT-qPCR compared to the t(11;14) KMS-12-PE cell lines. Points represent biological replicates, with error bars showing the mean ± SEM and significance determined using Welch’s two-sample t-tests on relative expression values. **(b)** Hierarchical clustering dendrogram from weighted gene co-expression network analysis (WGCNA) of malignant plasma cells, identifying co-expression modules used to infer nc209-associated gene programs. **(c)** Network visualization of the nc209-containing WGCNA module, M4, with nc209 highlighted in blue. Nodes are shaped by gene type, with inner nodes representing the top 10 hub genes in M4, outer nodes representing the remaining 30 genes of the top 40 hub genes, and edges indicating co-expression relationships. **(d)** Gene membership ranking for M4, showing genes ordered by module membership strength (kME) with nc209 among the highly connected genes in M4. Coding and noncoding genes are indicated by shape, and WGCNA module assignment is indicated by color. **(e)** Canonical WNT (cWnt) pathway enrichment in malignant plasma-cell pseudobulk profiles stratified by nc209 expression group, depicting increased cWnt pathway scores in nc209-high profiles with significance determined using a two-sided Wilcoxon rank-sum test with Benjamini-Hochberg correction. **(f)** Cell proliferation measured by OD absorbance at 450 nm following gapmer antisense oligonucleotide-mediated (ASO) knockdown (KD) of *ENSG00000310209* in the nc209-high t(4;14) myeloma cell lines H929 and KMS-18 relative to negative controls (NC) treated with transfection reagent alone. Points represent biological replicates, with error bars showing the mean ± SEM and significance determined using Welch’s two-sample t-tests. **(g)** Cell proliferation measured by OD absorbance at 450 nm following nc209 overexpression (OE) in the nc209-low MOLP-8 cell line, wherein nc209 OE increased proliferation relative to the empty vector (EV) control. Points represent biological replicates, with error bars showing the mean ± SEM and significance determined using Welch’s two-sample t-test. **(h)** Western blot analysis of β-catenin and c-MYC protein expression following nc209 knockdown in KMS-18 cells with GAPDH and histone H3 used as loading controls. nc209 knockdown reduced β-catenin and c-MYC protein expression, supporting an association between nc209 and WNT/β-catenin signaling. **(i)** Genomic view of the nc209 locus on chromosome 1q32.1, showing the *ENSG00000310209* transcript model, H3K4me3 and H3K27ac signals in bone marrow from the ENCODE4 database, and transcription factor binding in lymphoid and myeloma cell lines from the 2022 ReMap Chip-Seq Atlas. IRF4 ChIP-seq peaks were observed at the nc209 locus across lymphoid and myeloma cell lines. **(j)** Dumbbell plot comparing mean per-patient pseudobulk expression of genes from the Shaffer *et al.* myeloma IRF4 target gene set in nc209-low and nc209-high malignant plasma-cell profiles shows increased IRF4 activity in nc209-high versus nc209-low profiles. Significance determined using two-sided Wilcoxon rank-sum tests with Benjamini-Hochberg correction. **(k)** Relative IRF4 expression measured by RT-qPCR in the wild-type (WT) nc209-high KMS-18 cell line compared to the nc209-low KMS-12-PE EV control cell line. IRF4 expression was higher in KMS-18 than KMS-12-PE and increased following nc209 overexpression in KMS-12-PE cells. Points represent biological replicates, with error bars showing the mean ± SEM and significance determined using Welch’s two-sample t-tests. **(l)** Differential expression analysis of ligand-encoding genes comparing malignant plasma cells with nc209 expression in the upper quartile versus all other plasma cells with significance determined using a two-sided Wilcoxon rank-sum test with Benjamini-Hochberg correction. **(m)** Relative CCL5 expression measured by RT-qPCR in the nc209-high KMS-18 cell line compared to the nc209-low KMS-12-PE cell line, wherein CCL5 expression was increased following nc209 OE in KMS-12-PE cells and reduced following nc209 KD in KMS-18 cells. Points represent biological replicates, with error bars showing the mean ± SEM and significance determined using Welch’s two-sample t-tests. **(n)** Average per-sample pseudobulk expression of nc209 in malignant plasma cells across an independent single-cell RNA-seq cohort stratified by t(4;14) translocation status, illustrating higher nc209 expression in samples with t(4;14), with significance determined using a two-sided Wilcoxon rank-sum test with Benjamini-Hochberg correction. **(o)** Tumor-intrinsic cWnt pathway activity in the independent cohort, showing a trend toward increased activity in nc209-high tumor samples with significance determined using a two-sided Wilcoxon rank-sum test with Benjamini-Hochberg correction. Samples were stratified into high-and low-expression groups based on the median per-sample pseudobulk expression of nc209. **(p)** Average per-sample pseudobulk expression of CCL5 in the independent cohort stratified by nc209 expression group, displaying increased plasma cell CCL5 expression in nc209-high tumor samples with significance determined using a two-sided Wilcoxon rank-sum test with Benjamini-Hochberg correction.

We next reasoned that genes highly co-expressed with nc209 may be regulated by this lncRNA, either directly or indirectly, thereby providing insights into which oncogenic pathway(s) are associated with nc209. Weighted gene co-expression network analysis (WGCNA) of malignant plasma cells in the single-cell atlas identified 12 co-expression modules, with nc209 assigned to Module 4 (M4) (Figure 5b,c). Notably, M4 contained 10 of the 11 candidate ncRNAs derived from Cluster 12, suggesting that these transcripts may participate in a shared or closely related transcriptional program linked to the aggressive phenotype of this cluster. Examination of the 76 genes in M4 revealed several highly connected mRNAs with established roles in myeloma biology, including *CCND2* and *BAMBI*, implicating proliferative and Wnt-associated signaling programs (Figure 5d).^42,43^ Consistent with this observation, malignant plasma-cell pseudobulk profiles with high nc209 expression showed increased canonical Wnt signaling relative to nc209-low profiles (Figure 5e).

We then tested whether nc209 contributed functionally to myeloma cell proliferation. Gapmer antisense oligonucleotide (ASO)-mediated knockdown of nc209 in high-expressing t(4;14) myeloma cell lines reduced proliferation (Figure 5f), whereas overexpression of nc209 in the low-expressing t(11;14) cell line MOLP-8 increased proliferation (Figure 5g). In line with the Wnt-associated transcriptional signature, nc209 knockdown also reduced β-catenin and c-MYC protein expression in the KMS-18 cell line exhibiting high endogenous nc209 expression (Figure 5h). Together, these data support a role for nc209 in promoting myeloma cell fitness, at least in part through pathways linked to Wnt/β-catenin signaling.

To identify potential upstream regulators of nc209, we next examined publicly available chromatin immunoprecipitation sequencing (ChIP-seq) data for known binders of the *ENSG00000310209* locus. Interestingly, IRF4 was found to bind at the *ENSG00000310209* locus in several primary lymphocyte and myeloma cell lines, suggesting that nc209 may be regulated by IRF4 (Figure 5i). Supporting this possibility, nc209-high malignant plasma-cell profiles showed increased *IRF4* activity and expression (Figure 5j). In vitro, the nc209-high KMS-18 cell line also expressed higher *IRF4* than the nc209-low KMS-12-PE cell line, and overexpression of nc209 in KMS-12-PE cells increased *IRF4* expression (Figure 5k). These findings suggest a potential positive regulatory relationship between IRF4 and nc209.

Because immune dysregulation is a major determinant of MM progression,^21–23,44^ we next asked whether nc209-high myeloma cells expressed ligands that could influence the surrounding immune microenvironment. Differential expression analysis comparing malignant plasma cells in the top quartile of nc209 expression identified several upregulated ligands in nc209-high cells (log_2_fold-change > 1, adjusted *P* < 0.05; Figure 5l). Among the top candidates was *CCL5*, which was also present in the nc209-containing M4 module (Figure 5c). In vitro, the overexpression of nc209 increased CCL5 expression in KMS-12-PE cells, while the knockdown of nc209 expression significantly decreased CCL5 expression in KMS-18 cells (Figure 5m). These results nominate *CCL5* signaling as a potential link between nc209-high myeloma cells and immune microenvironmental remodeling, aligning with previous studies correlating CCL5 to the induction of immunosuppressive immune cells and poor treatment responses.^45–47^

Finally, we tested whether these tumor-intrinsic associations were reproduced in an independent clinical cohort with available raw bone marrow sequencing data and clinical treatment-response annotations, reported by Dhodapkar *et al*.^48^ We realigned the single-cell RNA-seq data from bone marrow aspirates collected from MM patients before (N = 6) and after (N = 16) CAR-T therapy and quantified noncoding gene expression by re-aligning the raw data to the integrated reference annotation (Figure S10). Consistent with our findings, samples from patients with t(4;14) myeloma showed higher nc209 expression than those without t(4;14) (Figure 5n). Moreover, nc209-high samples showed increased canonical Wnt and CCL5 signaling trends (Figure 5o,p), supporting the association between nc209, Wnt pathway activity, and chemokine signaling across independent datasets. Collectively, these findings identify nc209 as a previously uncharacterized t(4;14)-associated ncRNA linked to IRF4, Wnt/β-catenin signaling, CCL5 expression, and myeloma cell proliferation.

### Tumor-intrinsic nc209 expression is associated with CCL5 signaling and immune dysregulation

Because nc209-high malignant plasma cells showed increased *CCL5* expression in the discovery cohort, *in vitro* models, and an independent clinical dataset, we next asked whether the nc209-high tumor state was associated with remodeling of the surrounding immune microenvironment. For this analysis, we summarized nc209 expression in malignant plasma cells at the patient level using pseudobulk expression values from the single-cell atlas. Patients were then stratified into nc209-high and nc209-low groups using the median malignant plasma-cell pseudobulk nc209 expression as a cutoff, and immune subclusters were compared between groups to identify microenvironmental features associated with the nc209-high myeloma phenotype.

We first examined which immune subpopulations exhibited increased expression of the corresponding CCL5 receptors, CCR1, CCR3, and CCR5, in the nc209-high versus nc209-low tumors (Figure 6a). CCR5 receptor expression was broadly increased across lymphoid and myeloid subpopulations in nc209-high tumors, with the most prominent enrichment observed among CD8+ T effector populations. Given prior links between CCL5 signaling and immunosuppressive remodeling of the myeloma bone marrow microenvironment, we next compared CD8+ T effector profiles between nc209-high and nc209-low tumors (Figure 6b). Interestingly, CD8+ T effectors from nc209-high tumors showed increased expression of genes associated with inhibitory or dysfunctional T cell states, including the checkpoint molecule SIRPγ (*SIRPG*);^49^ H1 histones (*H1-4*, *H1-5*) that negatively regulate IL-2 and IFN-γ production;^50^ and TOX2 (*TOX2*), a regulator of T cell memory and exhaustion.^51,52^ These cells also showed increased expression of interferon-stimulated genes, including *IFI44L* and *GBP1*.^53^ Conversely, CD8+ T effectors from nc209-low tumors highly expressed genes encoding cytotoxic granzymes (*GZMH*, *GZMB*), cell motility proteins (*FRY*, *MYO3B*), and activation markers (*ITGB1*, *B2M*) as well as potentiators of CD8+ T effector function (*FGFBP2*, *ZNF683*, *CD226*).^54–56^ Together, these findings suggest that CD8+ T effector cells in nc209-high tumors adopt a transcriptional state characterized by interferon stimulation and reduced effector function.

**Figure 6.**
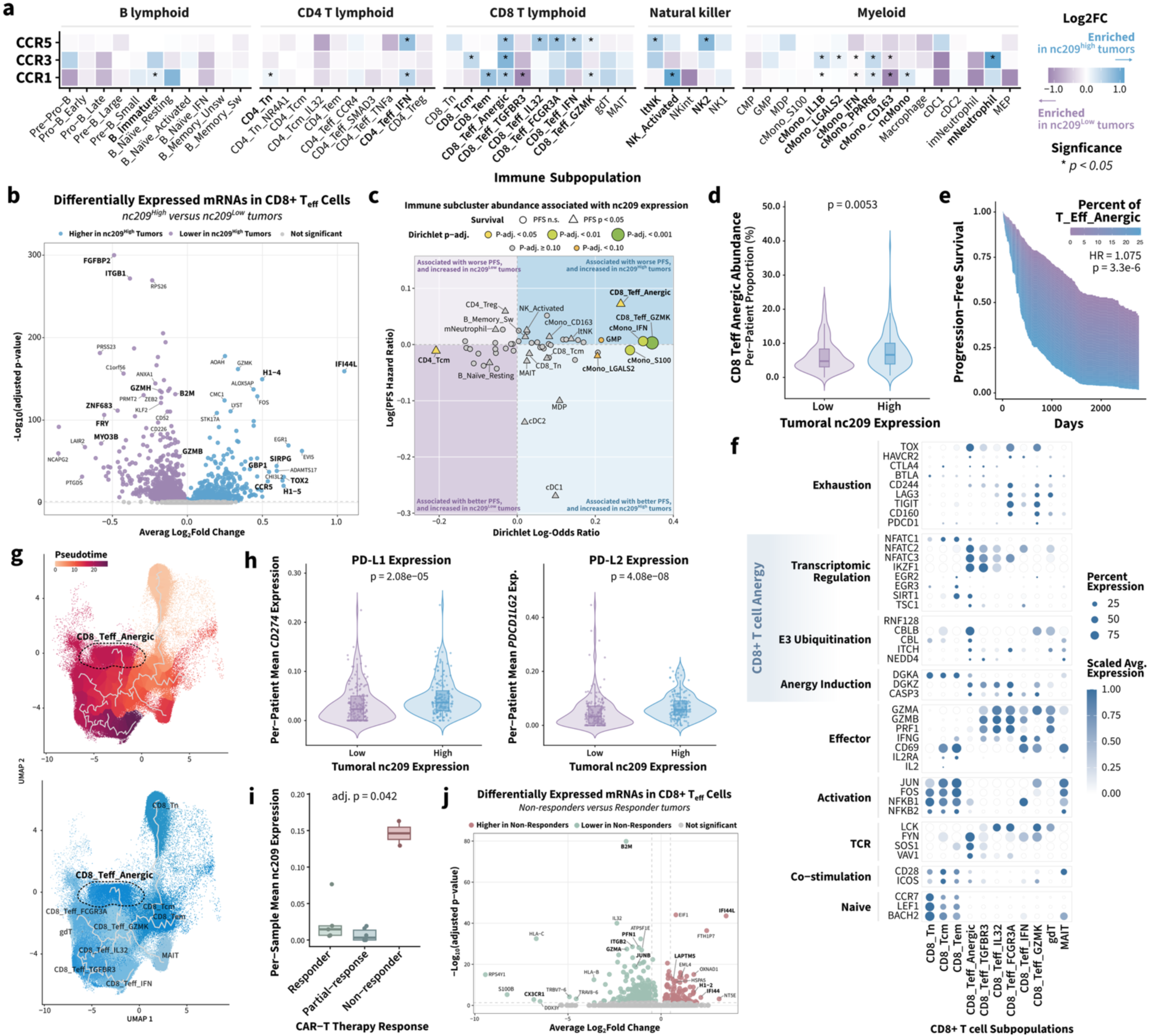
Tumor-intrinsic nc209 expression is associated with CCL5/CCR5 signaling, CD8+ T cell dysfunction, and impaired immunotherapy response. **(a)** Heatmap showing differential expression of the CCL5 receptors CCR1, CCR3, and CCR5 across immune subpopulations from patients stratified by malignant plasma-cell nc209 expression. Patients were classified as nc209-high or nc209-low using median malignant plasma-cell pseudobulk nc209 expression. Color indicates average log_2_fold-change, with blue representing enrichment in nc209-high tumors and purple representing enrichment in nc209-low tumors. Asterisks indicate significant differential expression as determined using a two-sided Wilcoxon rank-sum test. **(b)** Volcano plot showing differentially expressed mRNAs in CD8+ T effector cells from nc209-high (blue) versus nc209-low (purple) tumors with significance determined using a two-sided Wilcoxon rank-sum test with Bonferroni correction. **(c)** Scatter plot displaying the differential abundance of immune subclusters and the subcluster association with PFS; the x-axis shows the Dirichlet regression log-odds ratio for enrichment in nc209-high versus nc209-low tumors, and the y-axis shows the log hazard ratio for the Cox proportional hazards model of PFS based on the abundance of each subpopulation for all samples within the cohort. Each point represents an immune subpopulation. Point size and color indicate Dirichlet regression adjusted *P*-value (Wald test with Benjamini-Hochberg correction), while the point shape indicates survival significance. **(d)** Violin plots showing the increase in per-patient abundance of the CD8_Teff_Anergic subcluster in nc209-high versus nc209-low tumors with significance determined using a two-sided Wilcoxon rank-sum test. **(e)** Continuous Cox proportional hazards model showing the association between CD8_Teff_Anergic abundance (blue) using the per-patient proportion of the CD8_Teff_Anergic cluster within the CD8+ T cell compartment and PFS. Significance was determined using a two-sided Wald test. **(f)** Dot plot showing expression of canonical markers of exhaustion, transcriptional regulation, E3 ubiquitination, anergy induction, activation, TCR signaling, costimulation, and naïve T cell identity across CD8+ T cell subpopulations. Dot size indicates the percentage of cells within each subcluster expressing each gene, and color indicates scaled average expression. **(g)** Trajectory analysis showing pseudotime progression across states within the CD8+ T cell compartment. The highlighted CD8_Teff_Anergic population localizes near a terminal region of the CD8+ T cell differentiation continuum, suggesting this population is a terminally differentiated state. **(h)** Violin plots showing increased malignant plasma-cell pseudobulk expression of *CD274* and *PDCD1LG2*, encoding PD-L1 and PD-L2, respectively, in nc209-high versus nc209-low tumors with significance determined using a two-sided Wilcoxon rank-sum test. **(i)** Box plot showing per-sample mean of nc209 expression across CAR-T therapy response groups in the independent Dhodapkar *et al.* cohort, wherein nc209 expression was increased in non-responders compared with responders and partial responders. Significance was determined using a two-sided Kruskal-Wallis test with Bonferroni correction. **(j)** Volcano plot showing differentially expressed mRNAs in CD8+ T effector cells from CAR-T non-responders (green) versus responders (red), with significance determined using a two-sided Wilcoxon rank-sum test with Bonferroni correction.

Consistent with this pattern, the CD8_Teff_Anergic subcluster, which showed increased CCR5 expression, was significantly enriched in nc209-high tumors (Figure 6c,d) and associated with worse PFS (Figure 6e). To further characterize this population, we examined canonical markers of T cell activation, exhaustion, anergy, costimulation, and effector function across CD8+ T cell subclusters (Figure 6f). The CD8_Teff_Anergic population expressed the co-stimulatory molecules *CD28* and *ICOS,* as well as the TCR signaling components *FYN*, *SOS1*, and *VAV1*. However, this population showed limited expression of downstream activation (e.g., *JUN*, *FOS, NFKB1, NFKB2*) and effector (e.g., *GZMA, GZMB, PRF1, IFNG*) genes, while expressing established anergy-associated regulators, including *DGKZ*, *CASP3*, and *CBLB*,^57–59^ and the activation regulator, *TOX*. This profile is consistent with an anergic-like CD8+ T cell state marked by preserved TCR stimulation but attenuated downstream activation and effector function. Trajectory analysis further placed the CD8_Teff_Anergic population as a terminal state on the CD8+ T cell differentiation continuum, supporting its interpretation as a terminal dysfunctional effector state (Figure 6g).

Because CD8+ T cell anergy can be induced by multiple tumor microenvironmental mechanisms, including inhibitory checkpoint signaling, we next evaluated their expression across malignant and immune compartments. Notably, malignant plasma cells from nc209-high tumors showed significantly increased expression of *CD274* and *PDCD1LG2*, encoding PD-L1 and PD-L2, respectively (Figure 6h). These findings suggest that nc209-high malignant plasma cells may contribute to pro-tumoral remodeling of the immune microenvironment through both chemokine-mediated signaling and increased expression of inhibitory checkpoint ligands.

Lastly, to extend these findings to our independent validation cohort, we examined whether the nc209-associated tumor and immune features identified above were also linked to clinical treatment response. We began by comparing nc209 expression across response groups and found that nc209 was increased in samples from non-responders relative to responders and partial responders (Figure 6i), suggesting that the nc209-associated myeloma phenotype may be linked to immunotherapy response. We next compared CD8+ T effector transcriptional profiles between non-responders and responders to determine whether the immune features associated with nc209-high tumors in the discovery cohort were also evident in this independent clinical context (Figure 6j). Notably, we observed that non-responders exhibited decreased expression of activation (*JUNB*, *ITGB2, B2M*) and cytotoxic effector (*PFN1*, *GZMA, CX3CR1*) markers,^54^ with increased expression of interferon-induced genes (*IFI44L, IFI44*) and negative regulators of T cell activation (*LAPTM5*, *H1-2*),^50,60^ consistent with the interferon-stimulated anergic-like CD8+ T effector profile observed in nc209-high tumors from our discovery cohort (Figure 6b,f). Collectively, these findings link the nc209-high tumor state to changes in the immune microenvironment, including CCL5-CCR5 signaling, increased immune checkpoint expression, outcome-associated CD8+ T cell dysfunction, and poorer CAR-T response.

## Discussion

In this study, we developed and benchmarked an integrated coding-noncoding reference annotation that enables concurrent interrogation of coding and noncoding transcription from standard RNA-sequencing data. By systematically resolving conflicts between genomic coordinates of overlapping coding and noncoding transcripts, the resulting reference preserved the established coding gene space while substantially expanding the accessible noncoding transcriptome. Alignment to the resulting integrated genome annotation reduced ambiguous read assignment relative to direct annotation concatenation, increased ncRNA detection without reducing coding gene capture, and maintained the major cellular architecture recovered by conventional single-cell analysis. Application to large bulk and single-cell multiple myeloma cohorts further demonstrated that the newly accessible ncRNAs were not restricted to low-information transcriptional background. Instead, they contributed to highly variable features, cell-type-specific profiles, clinically relevant immune states, and tumor phenotypes associated with patient outcomes. Together, these findings establish our integrated reference annotation as a valuable tool for accessing the expansive noncoding genome from existing and future transcriptomic data, fostering biomarker discovery, mechanistic investigation, and therapeutic development.

Previous annotation efforts have made essential but incongruous contributions to defining the human transcriptome; curated references such as GENCODE prioritize transcript support and annotation confidence, whereas noncoding resources such as LncBook aim to capture the largely uncharacterized noncoding genome. However, combining these independently constructed annotations to obtain a more comprehensive survey of the transcriptome cannot be achieved by simple concatenation due to genomic overlap among coding, antisense, and sense-overlapping noncoding transcripts. The resulting conflicts in genomic coordinates produce ambiguous reads during sequencing alignment, which are subsequently discarded, thereby limiting transcript capture and complicating downstream analysis. Our results demonstrate that resolving these conflicts through systematic integration of the coding and noncoding genome annotations is critical. The integrated reference retained 175,832 genes, including 138,296 noncoding genes, while limiting trimming to a minority of ncRNA transcripts. This strategy therefore balances preserving confidence in established coding annotations and expanding access to the noncoding transcriptome.

The benchmarking analyses further indicate that expanded ncRNA detection can be incorporated without destabilizing standard single-cell workflows. The integrated and GENCODE-aligned datasets retained highly concordant cell recovery, quality metrics, compartment abundance, and broad cell-type assignments, while the integrated reference increased the representation of ncRNAs among highly variable genes and differentially expressed features. This observation is consistent with prior work showing that lncRNAs frequently display greater tissue and cell-state specificity than protein-coding genes. In the myeloma bone marrow atlas, noncoding genes accounted for a substantial proportion of subcluster-defining transcripts and distinguished outcome-associated immune populations. Thus, excluding broader ncRNA annotations may preferentially remove features that encode cellular identity and disease-associated features, which are central to single-cell cancer studies.

Leveraging our integrated genome annotation, we generated a coding-noncoding atlas of the myeloma bone marrow microenvironment to identify novel ncRNA candidates related to poor patient outcomes in high-risk myeloma. Extending prior transcriptomic classifications of myeloma,^23,36^ showing that expression-defined patient groups capture biological and prognostic heterogeneity beyond individual cytogenetic lesions, we defined 15 patient clusters based on their coding and noncoding gene expression profiles, two of which were significantly associated with poor outcomes and enriched for t(4;14) and 1q21+ abnormalities. Joint coding-noncoding analysis identified 19 candidate ncRNAs that were highly specific to these high-risk patient groups, outcome-associated, concordant across modalities, and preferentially expressed in malignant plasma cells. The recovery of *LINC01287*, a transcript previously implicated in several solid tumors but not yet investigated in the context of myeloma, as our top candidate ncRNA, provided support for the ability of our integrated analytical approach to identify conserved cancer-associated ncRNA relationships.

Moreover, among the 19 candidate ncRNAs, 16 had not been previously characterized, underscoring the ability of our approach to provide access to the vastly understudied noncoding genome for identifying novel biomarkers, mechanistic insights, and therapeutic targets. To probe these novel candidate ncRNAs, we characterized *ENSG00000310209,* which we labeled nc209, and revealed that it is associated with a high-risk t(4;14)-enriched tumor state, proliferative fitness, Wnt pathway activation, CCL5 expression, and immune dysfunction, providing convergent evidence that the workflow can nominate previously uncharacterized transcripts with biological and clinical relevance. These findings do not establish nc209 as the dominant regulator of these programs, nor imply that all newly detected ncRNAs are functional. Rather, they demonstrate that annotation expansion, cohort-scale prioritization, cellular localization, clinical association, and experimental perturbation can distinguish plausible disease-associated candidates from the larger background of the incompletely characterized noncoding transcriptome.

The broader value of this approach lies in its compatibility with the large body of RNA-sequencing data already generated across cancer and other diseases. Large consortia, institutional biobanks, clinical trials, and single-cell atlases contain raw sequencing reads that are commonly analyzed using references optimized for established genes. Re-alignment to an integrated coding-noncoding annotation could expose additional disease-, lineage-, and treatment-associated signals without requiring new sample acquisition. Because the approach preserves simultaneous analysis of coding pathways and ncRNA expression, candidate noncoding genes can be interpreted directly in relation to oncogenic signaling, cellular composition, treatment response, and survival. This is particularly relevant for diseases in which rare cellular states or transcriptional subtypes influence outcome, but the same principle could extend beyond cancer to developmental, inflammatory, cardiovascular, and neurological datasets in which noncoding regulation is strongly context dependent.

Several limitations should be considered. First, the integrated reference remains dependent on the accuracy and completeness of its source annotations. LncBook includes predicted and variably supported transcripts, and some of these annotated noncoding transcripts may represent incomplete isoforms, unstable transcription, or transcriptional noise. Conversely, prioritizing coding annotations and trimming overlapping ncRNAs may remove functional sequences or reduce quantification of biologically relevant sense and antisense ncRNAs. Second, short-read bulk and 3′ single-cell RNA sequencing cannot reliably resolve full-length transcript structures and may under-detect low-abundance or non-polyadenylated ncRNAs, including microRNAs (miRNAs). Third, our benchmarking analysis revealed minor discordance between the annotations of several immune subpopulations in myeloma bone marrow (Figure 2h), which could not be adjudicated against cell-specific ground truth. Finally, the function of nc209 was inferred from *in silico* expression, co-expression, and survival analyses. The *in vitro* nc209 experiments provide preliminary functional validation to highlight the utility of our integrated analytical approach for identifying biologically relevant candidate ncRNA, but causal links among tumor ncRNA expression, immune remodeling, and therapeutic response require further testing in primary samples and *in vivo* models.

Collectively, this work presents an integrated coding-noncoding human reference genome annotation and analytical strategy for recovering comprehensive, clinically informative transcriptome profiles from conventional sequencing data. The application to multiple myeloma demonstrates that expanded annotation can preserve established biological structure while revealing noncoding cell-state markers, prognostic candidates, and previously uncharacterized ncRNAs that are inaccessible to coding-centered approaches. More broadly, our integrated genome annotation opens the door to mining the expansive noncoding genome using existing sequencing datasets that remain constrained by the genome references used to interpret them. Systematic reanalysis with comprehensive, ambiguity-resolved annotations may therefore unlock a substantial layer of disease biology already present within current genomic resources, yielding novel biomarkers and therapeutic targets, as well as mechanistic insights into pathogenesis and treatment responses.

## Methods

### Development of the integrated coding-noncoding reference genome annotation

We developed a five-step computational pipeline to integrate LncBook v2.1 with GENCODE v47 while preserving the original coding annotation.^13,15^ In the first step, the input annotations were standardized and validated. In the second step, noncoding transcripts were classified according to their genomic overlap with coding genes. Non-overlapping transcripts were retained without modification, whereas sense-and antisense-overlapping ncRNA transcripts were processed using strand-aware subtraction to remove regions shared with coding features in the third step. In the fourth step, an additional 250-bp buffer was removed from each trimmed boundary to reduce ambiguous read assignment near coding-noncoding junctions, and transcripts with insufficient remaining sequence were discarded. In the final step, the retained noncoding transcripts were consolidated, validated, and combined with the preserved coding annotation to generate the final integrated coding-noncoding reference genome annotation. 10x Genomics Cell Ranger (v8.0.1)^61^ and STAR (v2.7.11b)^62^ reference indices were then generated using the GRCh38.p14 genome assembly and the integrated annotation for alignment of single-cell and bulk RNA-sequencing datasets, respectively. A detailed description of each step of the generated pipeline is available in Supplemental Methods S1.

### Benchmarking the integrated coding-noncoding reference genome annotation

The integrated annotation was benchmarked using 48 single-cell RNA-sequencing samples randomly selected across four sequencing batches. Each sample was independently processed against three reference annotations: GENCODE v47, a direct concatenation of GENCODE v47 and LncBook v2.1 without resolution of overlapping loci, and the integrated GENCODE v47-LncBook v2.1 annotation. A detailed description of the benchmarking analyses is available in Supplemental Methods S2.

To briefly summarize, identical Cell Ranger and quality-control procedures were applied to each alignment condition. First, cells with greater than 1,000 UMIs, more than 200 detected features, and less than 20% mitochondrial transcript abundance were retained. Genes with greater than 10 total counts, detected in more than 10 cells, and expressed in more than 20% of samples were retained. Reference performance was evaluated by comparing ambiguous read alignment, recovery of coding and noncoding features, per-cell UMI and feature counts, and cell-barcode retention. To determine whether the expanded annotation affected downstream biological interpretation, the GENCODE-and integrated-aligned datasets were generated using Seurat (v5.4.0),^63^ independently normalized, batch-corrected using Harmony (v.2.0.5),^64^ clustered, and annotated using identical parameters. Major cellular compartments and immune subpopulations were identified using canonical markers, and annotation concordance and population-specific differential expression were compared between references.

### Single-cell RNA-sequencing dataset processing and analyses

Single-cell RNA-sequencing samples from the MMRF CoMMpass study (N = 478) were aligned to the integrated reference and filtered to retain cells with more than 1,000 UMIs, more than 200 detected features, and less than 20% mitochondrial transcript abundance. Genes were retained when they had more than 10 total counts, were detected in more than 10 cells, and were expressed in more than 20% of samples. The final count matrix was converted into a Seurat v5 object using BPCells (v0.3.1) to support memory-efficient analysis, followed by count normalization and scaling of the 3,000 most variable features. A representative 10% sketch of the complete dataset was used for dimensionality reduction, Harmony batch correction, graph-based clustering, and UMAP generation. Cluster assignments were subsequently projected to the complete dataset. Major cellular compartments were annotated using canonical markers and previously established MMRF Immune Atlas annotations,^23^ with murine contaminants, putative doublets, and low-quality populations excluded. B lymphoid, myeloid, and NK and T lymphoid compartments were reclustered independently to resolve immune subpopulations. Subtype-specific markers and ncRNAs were identified, and associations between baseline immune composition and progression-free survival were evaluated using patient-level multivariable Cox proportional-hazards adjusted for sample processing study site and batch. A more detailed description of these methodologies, as well as downstream analyses of nc209 function and the immune compartment, is available in Supplemental Methods S3.

### Bulk RNA-sequencing dataset processing and analyses

Gene-level counts from 942 CD138-positive bulk RNA-sequencing samples from the MMRF CoMMpass study were generated by alignment to the integrated reference and combined into a gene-by-sample count matrix. The 776 baseline samples were used for all downstream analyses. Lowly detected genes and genes with extreme total count distributions were removed, after which counts were normalized and variance-stabilized using DESeq2.^65^ Patients were grouped according to their coding-noncoding tumor-expression profiles using consensus partitioning around medoids clustering of the top 10% most variable genes, yielding 15 patient clusters. Associations between cluster membership and progression-free survival were examined using multivariable Cox proportional-hazards models adjusted for autologous stem-cell transplantation, race, and sex. Cluster-specific ncRNAs were filtered by differential expression magnitude, statistical significance, cluster specificity, and association with progression-free survival to yield an initial list of candidates. These candidate ncRNAs were then evaluated in the single-cell atlas to confirm directionally concordant expression in malignant plasma cells and enrichment within the plasma-cell compartment, producing a final set of 19 candidates. These candidates were ranked using a weighted composite score incorporating the absolute cluster-specific log_2_fold-change, differential-expression significance, magnitude of the progression-free-survival hazard ratio, and Cox regression significance. For a more detailed description of the bulk RNA-sequencing dataset processing, patient clustering, and ncRNA candidate selection, please see Supplemental Methods S4.

### In vitro characterization of nc209

A detailed description of the in vitro methodologies is available in Supplemental Methods S5. To briefly summarize, multiple myeloma cell lines were cultured under standard conditions and used to evaluate the functional consequences of altering nc209 expression. Loss-of-function experiments were performed by transfecting cells with three independent nc209-targeting GapmeR antisense oligonucleotides, with knockdown confirmed by RT-qPCR. For gain-of-function experiments, the nc209 sequence was cloned into an expression vector, and successfully transfected GFP-positive cells were isolated by flow cytometry. Cell proliferation and viability were measured 72 hours after transfection using the Dojindo Cell-Counting Kit 8 (Dojindo, Cat: CK04-11) assay across replicate wells. Changes in protein expression associated with nc209 were evaluated by western blotting, while changes in RNA expression were measured by SYBR Green-based RT-qPCR. Relative expression was calculated using the comparative threshold-cycle method and normalized to the appropriate housekeeping gene and experimental control.

## Supporting information

High Resolution Figures

## List of abbreviations

ASO: Antisense oligonucleotide
BMME: Bone marrow microenvironment
cDC: Classical dendritic cell
ChIP-seq: Chromatin immunoprecipitation sequencing
cWnt: Canonical Wnt
DEG: Differentially expressed gene
EV: Empty vector
GTF: Gene transfer format
IME: Immune microenvironment
ISG: Interferon-stimulated gene
KD: Knockdown
kME: Eigengene-based connectivity (module membership)
lncRNA: Long noncoding ribonucleic acid
MAIT: Mucosal-associated invariant T cell
MM: Multiple myeloma
MMRF: Multiple Myeloma Research Foundation
mRNA: Messenger ribonucleic acid
NC: Negative control
ncRNA: Noncoding ribonucleic acid
OD: Optical density
OE: Overexpression
RNA: Ribonucleic acid
WGCNA: Weighted-gene co-expression network analysis
WT: Wild type

## Declarations

### Ethics approval and participant consent

Samples used to generate the coding-noncoding single-cell and bulk RNA-sequencing datasets were originally collected through the MMRF CoMMpass study (<u>NCT01454297</u>). The CoMMpass protocol was approved by the institutional review board or independent ethics committee at each participating center, and all participants provided written informed consent for the collection, genomic analysis, and research use of their samples and clinical information. The study was conducted in accordance with the Declaration of Helsinki and applicable ethical and regulatory requirements. The independent validation analysis used previously published, deidentified single-cell RNA-sequencing data obtained from the Gene Expression Omnibus under accession <u>GSE210079</u>.

### Data availability

The raw single-cell and bulk RNA-sequencing data and associated clinical information obtained from the Multiple Myeloma Research Foundation (MMRF) are available under controlled access through the MMRF Virtual Lab (VLAB; https://mmrfvirtuallab.org/) data sharing platform. Raw single-cell RNA-sequencing data from the independent validation cohort are publicly available through the Gene Expression Omnibus under accession <u>GSE210079</u>. The integrated human reference genome generated in this study will be available through Zenodo upon publication (<u>10.5281/zenodo.21738380</u>).

### Code availability

All scripts generated for the presented analyses and figures are available on the MMRF Immune Atlas Consortium GitHub (https://github.com/theMMRF/MMRF_ImmuneAtlas).

### Declaration of interests

S.G. reports other research funding from Boehringer-Ingelheim, Bristol-Myers Squibb, Celgene, Genentech, Regeneron and Takeda and consulting from Taiho Pharmaceuticals, not related to this study. S.K. declares research funding for clinical trials to the institution from Abbvie, Amgen, Allogene, BMS, Carsgen, GSK, Janssen, Roche-Genentech, Takeda and Regeneron, as well as consulting or advisory board participation (with no personal payments) for Abbvie, BMS, Janssen, Roche-Genentech, Takeda, Pfizer, Loxo Oncology, K36, Sanofi, ArcellX and Beigene. D.A. declares grants from MMRF, CTN (National Heart, Lung and Blood Institute), Celgene, Pharmacyclics and Kite Pharma, as well as other support from Juno, Partners TX, Karyopharm, BMS, Aviv MedTech, Takeda, Legend Bio Tech, Chugai, Caribou Biosciences, Janssen, Parexel, Sanofi and Kowa. D.A. also has a patent (PCT/US2021/059199) pending. I.S.V. reports grants from NCI, National Heart, Lung, and Blood Institute, National Institute of Diabetes and Digestive and Kidney Diseases, Harvard Stem Cell Institute and consulting for Mosaic, AlphaSights, NextRNA and Guidepoint Global outside of the submitted work. The other authors declare no competing interests.

## Author information

### Author contributions

M.E.M. and D.O. contributed equally to this work.

## Consortia

### Immune Atlas Consortium

William C. Pilcher, Edgar Gonzalez-Kozlova, Dimitra Karagkouni, Chaitanya R. Acharya, Marina E. Michaud, Yizhe Song, Julia T. Wang, Sarthak Satpathy, Yuling Ma, Darwin D’Souza, Reyka G. Jayasinghe, Denis Ohlstrom, Katherine E. Ferguson, Giulia Cheloni, Mojtaba Bakhtiari, Kai Nie, Jennifer A. Foltz, Isabella Saldarriaga, Rania Alaaeldin, Rachel Chen, Mark A. Fiala, Junia Vieira Dos Santos, I-ling Chiang, Igor Figueiredo, Julie Fortier, Michael Slade, Stephen T. Oh, Michael P. Rettig, Emilie Anderson, Ying Li, Surendra Dasari, Michael A. Strausbauch, Travis Dawson, Brian H. Lee, Geoffrey Kelly, Laura Walker, Nicolas F. Fernandez, John Leech, Jarod Morgenroth-Rebin, Krista Angeliadis, Matthew A. Wyczalkowski, Song Cao, Omar Ibrahim, Roderick Lin, Todd A. Fehniger, Andrew Houston, Emir Radkevich, Adeeb H. Rahman, Zhihong Chen, Alessandro Lagana, John F. DiPersio, Jacalyn Rosenblatt, Seunghee Kim-Schulze, Sagar Lonial, Shaji Kumar, Swati S. Bhasin, Taxiarchis Kourelis, Madhav V. Dhodapkar, Ravi Vij, David Avigan, Hearn J. Cho, George Mulligan, Li Ding, Sacha Gnjatic, Ioannis S. Vlachos, and Manoj Bhasin

## Acknowledgements

Workflow schematics used in the figures were created using BioRender.com (https://BioRender.com/lpov86o). This work was made possible by funding support from the Multiple Myeloma Research Foundation (M.B., S.G., S.M., D.A. and I.S.V.), Myeloma Solutions Fund (M.B., D.O., M.E.M. and W.C.P.), and the Paula C. and Rodger O. Riney Foundation (S.M. and L.B.)

