## Supplementary figures and images for "Integrated coding-noncoding genome annotation expands single-cell transcriptomic discovery and identifies clinically relevant noncoding RNAs in multiple myeloma"

### High Resolution Figures

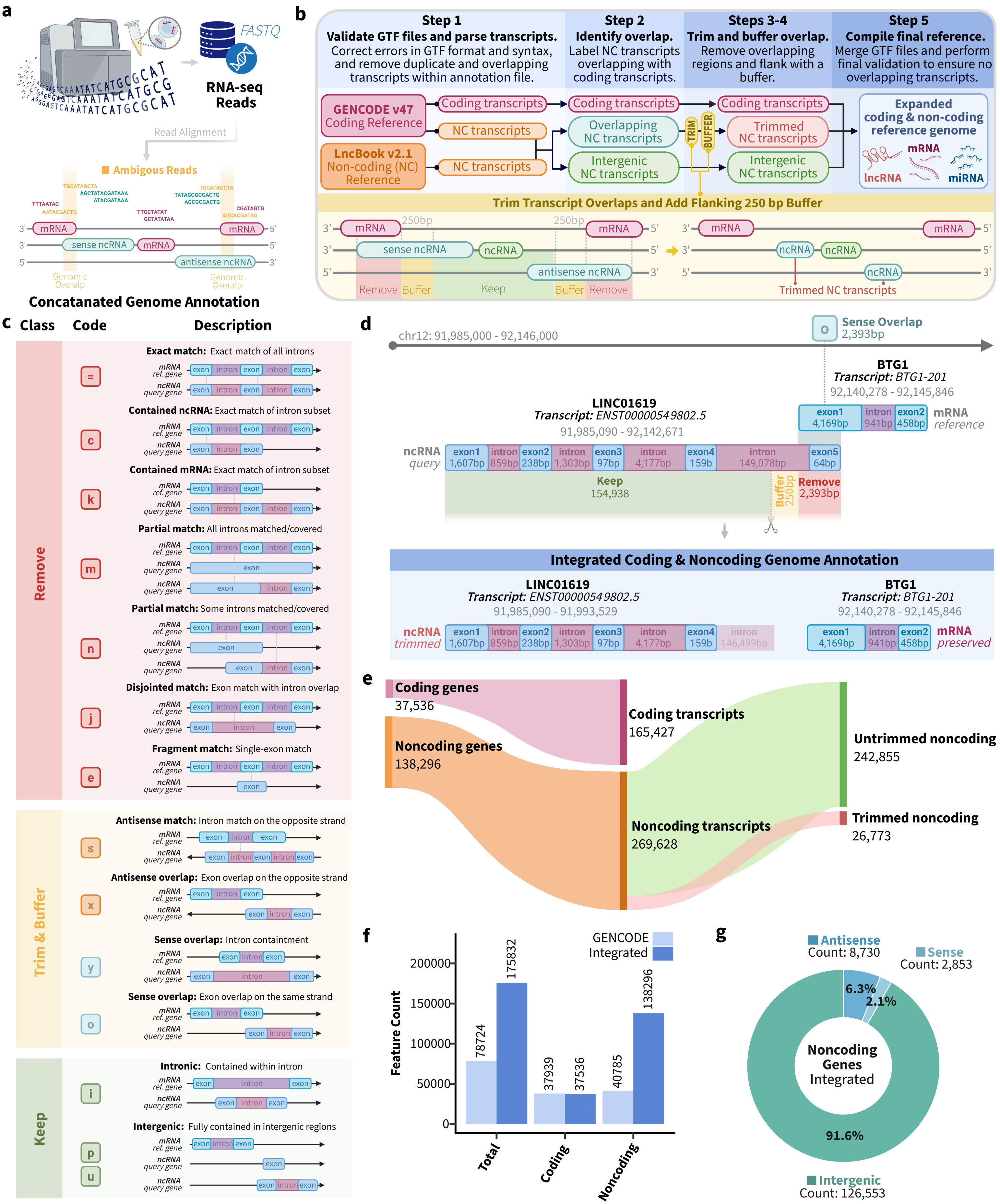

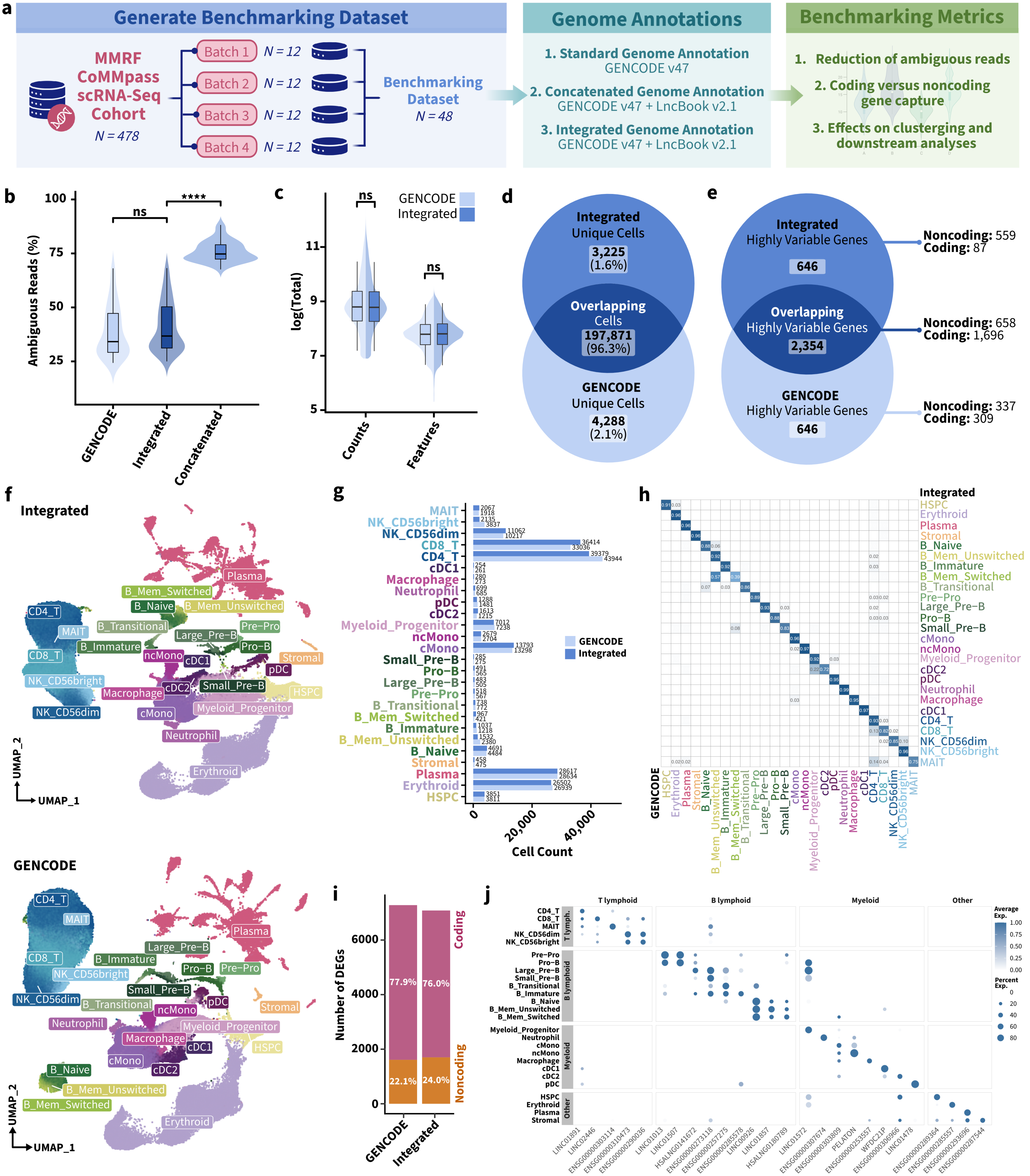

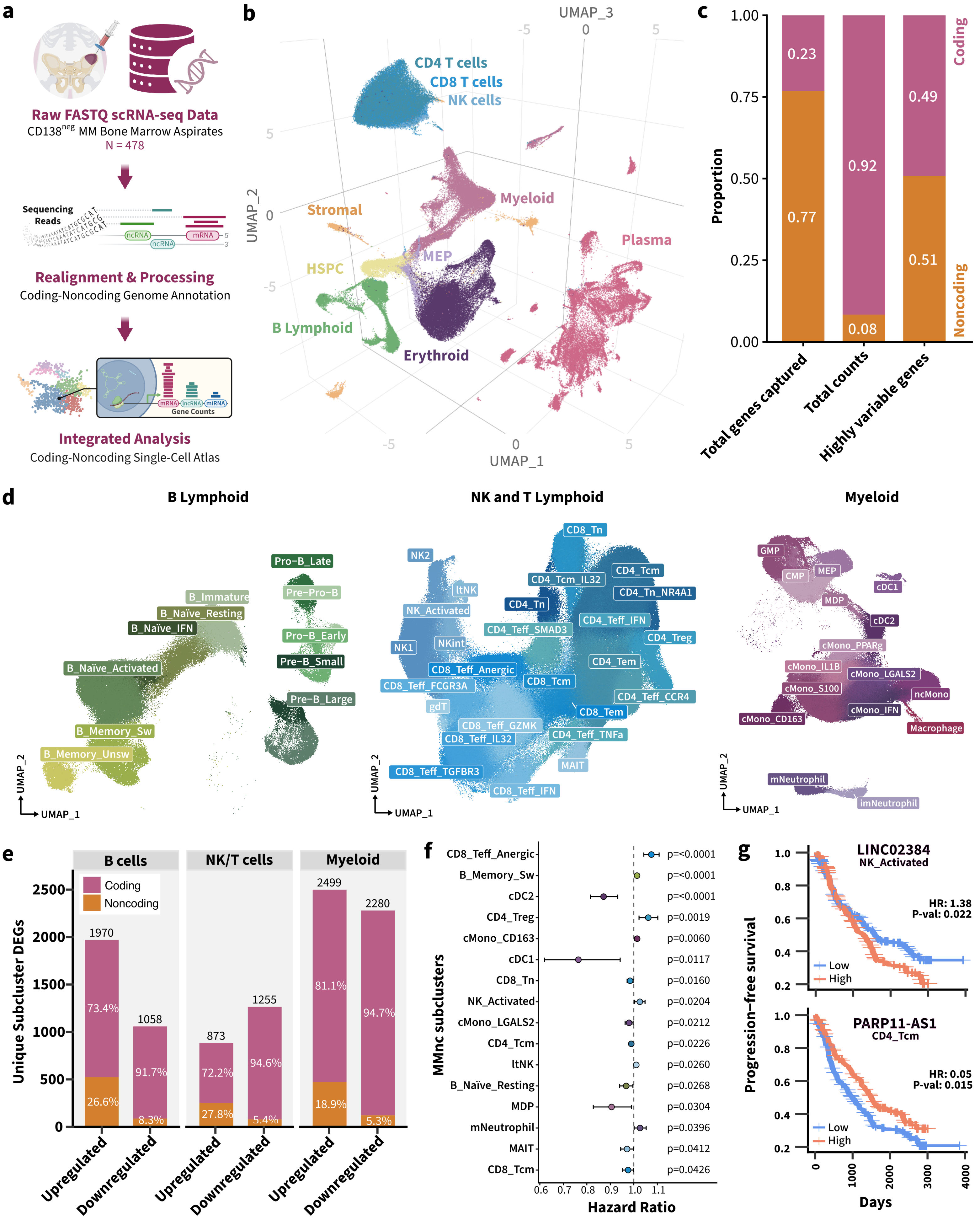

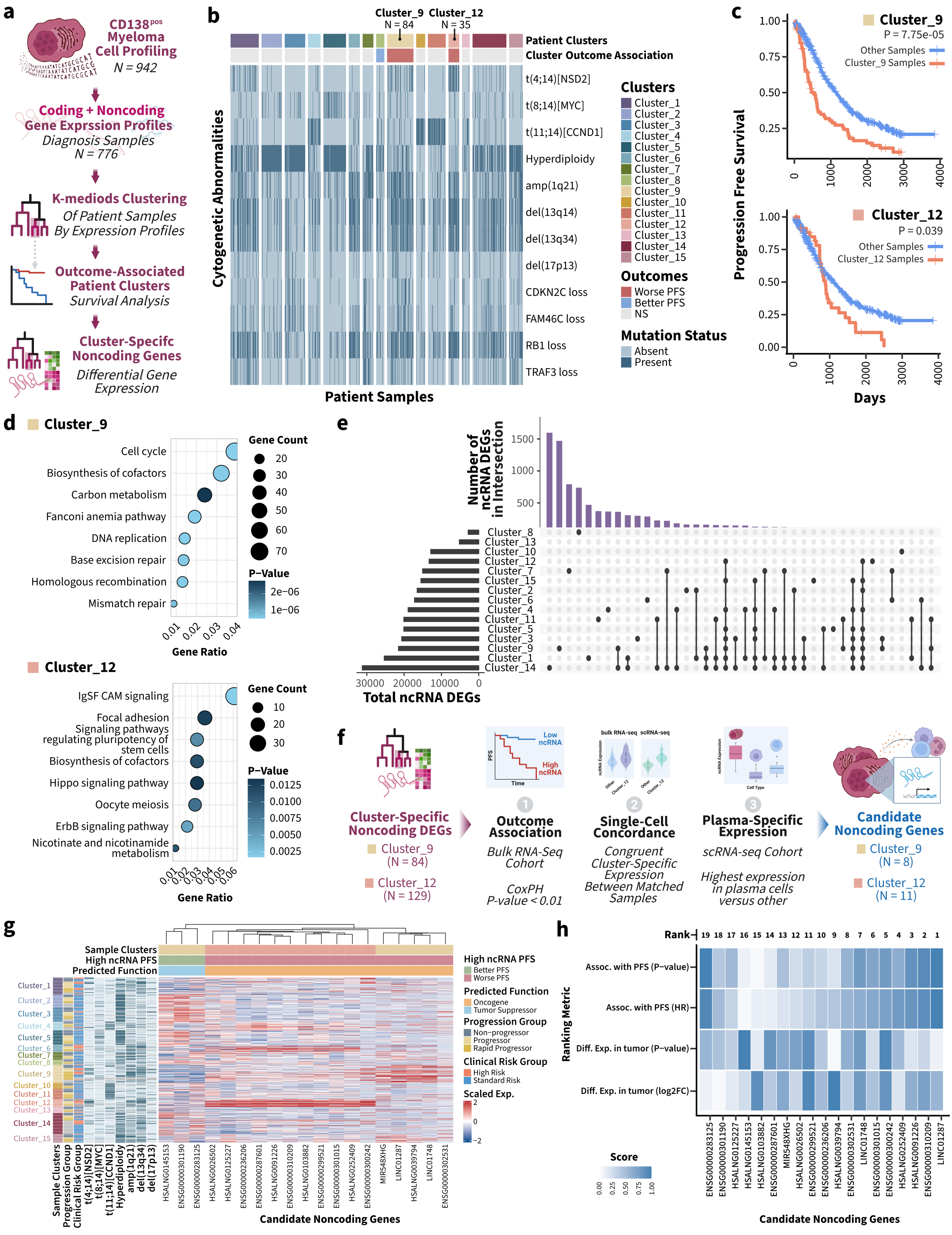

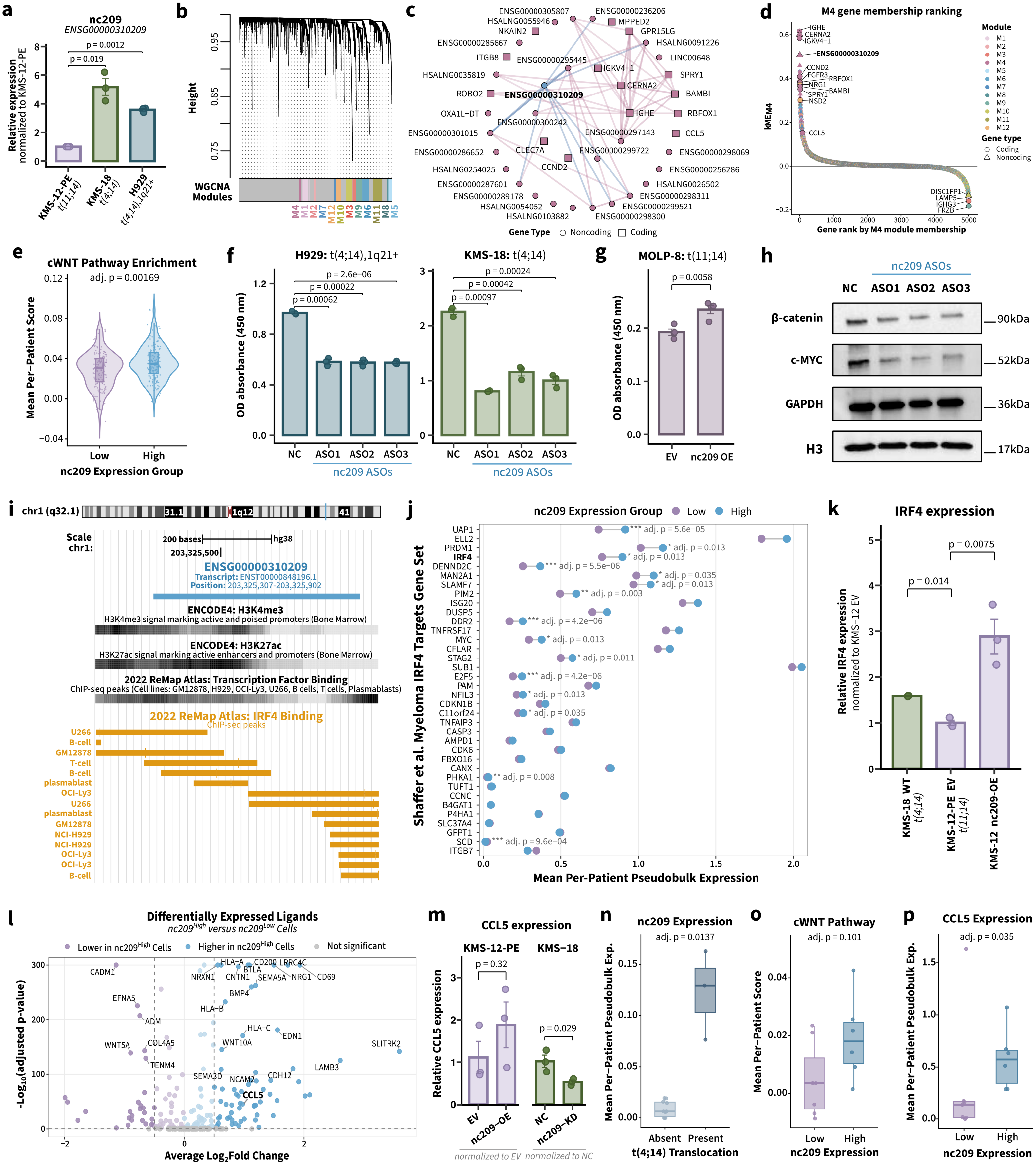

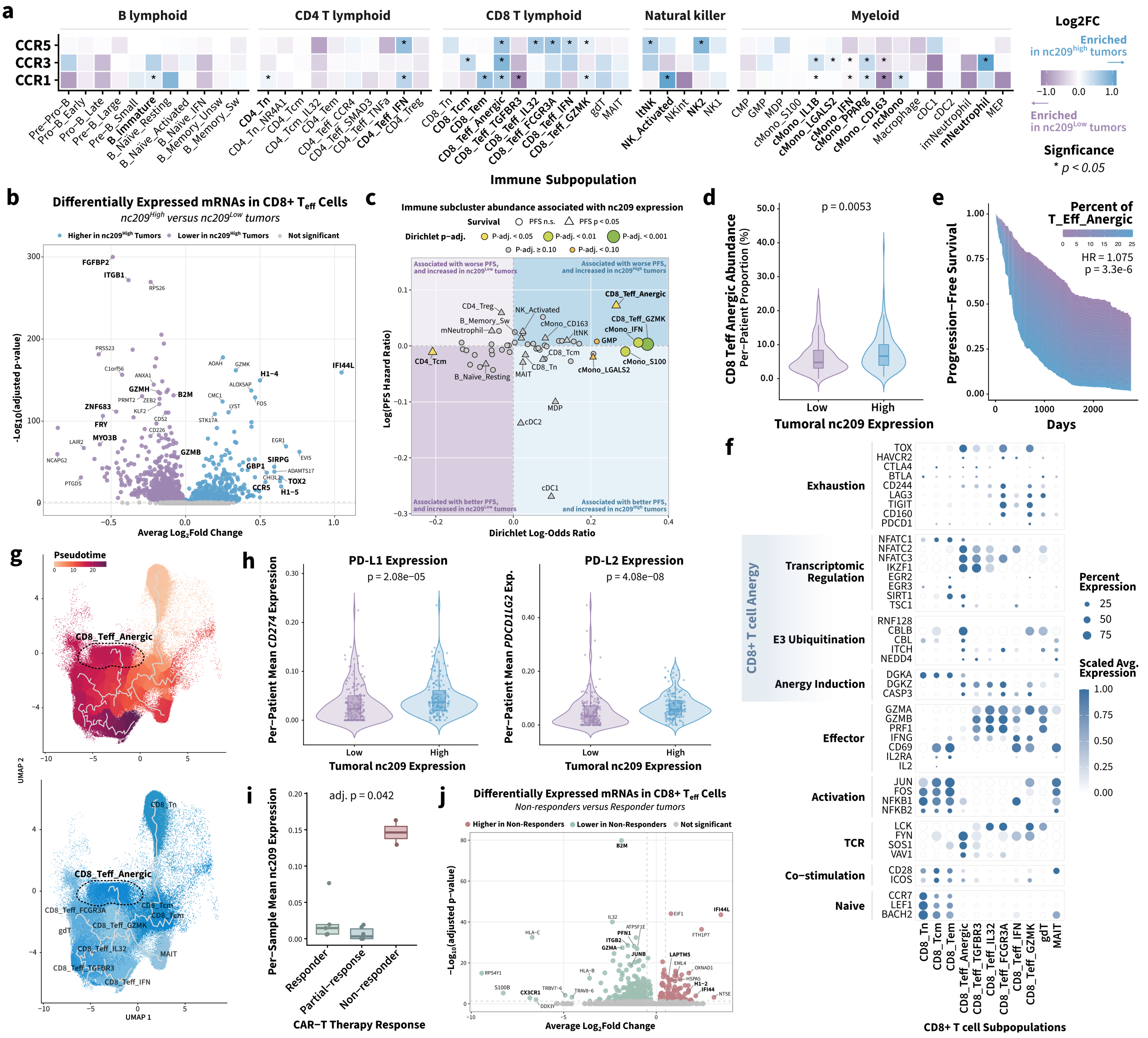
