## Supplemental Materials for "Integrated coding-noncoding genome annotation expands single-cell transcriptomic discovery and identifies clinically relevant noncoding RNAs in multiple myeloma"

### Table of Contents

#### Supplemental Tables

**Table S1.** GapmeR sequences used for nc209 knockdown………………………………………… 3

**Table S2.** PCR primer sequences……………………………………………………………….. 3

**Table S3.** Enriched pathways among samples in Cluster 9………………………………………. 4

**Table S4.** Enriched pathways among samples in Cluster 12…………………………………….. 5

#### Supplemental Figures

**Figure S1.** Quality control metrics assessing the benchmarking datasets……………………………. 6

**Figure S2.** Annotation of the complete coding-noncoding single-cell atlas…….…………………….. 7

**Figure S3.** Annotation of immune compartments within the coding-noncoding single-cell atlas…… 8

**Figure S4.** Annotation of immune subclusters within the coding-noncoding single-cell atlas……… 9

**Figure S5.** Association of immune subclusters with progression-free survival………………………. 10

**Figure S6.** LINC02384 association with interferon-gamma expression………………………………. 11

**Figure S7.** Correlation between tumor burden and single-cell plasma counts……………………….. 12

**Figure S8.** Patient cluster comparison and non-malignant expression of candidate ncRNAs……… 13

**Figure S9.** Expression of *LINC01287*-linked mRNA targets…………………………………………….. 15

**Figure S10.** External validation dataset annotation…………………………………………………… 16

#### Supplemental Methods

**Method S1.** Development of an integrated coding-noncoding reference genome annotation………. 17

**Method S2.** Benchmarking of the integrated reference genome annotation…………………………... 18

**Method S3.** Single-cell RNA-sequencing dataset processing and analyses………………………… 19

**Method S4.** Bulk RNA-sequencing dataset processing and analyses…………………………………. 23

**Method S5.** In vitro quantification of gene expression……………………………………………………. 24

**References**……………………………………………………………………………………………. 27

#### Supplemental Tables

**Table S1.** GapmeR sequences used for nc209 knockdown.

| ID | Target | Sequence | Vendor |
|---|---|---|---|
| **ASO1** | *ENSG00000310209.1* | GGAGCTGTGTGACCAAAGGA | Integrated DNA Technologies |
| **ASO2** | *ENSG00000310209.1* | ATCCCAGTGCAGGAGCCAGA | Integrated DNA Technologies |
| **ASO3** | *ENSG00000310209.1* | GACACCAGATCCACCTTACA | Integrated DNA Technologies |

**Table S2.** PCR primer sequences.

| Target | Fwd/Rev | Primer Sequence | Vendor |
|---|---|---|---|
| ***β-actin*** | Forward | GAATCAATGCAAGTTCGGTTCC | Integrated DNA Technologies |
| | Reverse | TCATCTCCGCTATTAGCTCCG | Integrated DNA Technologies |
| ***18S*** | Forward | ACCCGTTGAACCCCATTCGTGA | Integrated DNA Technologies |
| | Reverse | GCCTCACTAAACCATCCAATCGG | Integrated DNA Technologies |
| ***CCL5*** | Forward | CCAGCAGTCGTCTTTGTCAC | Integrated DNA Technologies |
| | Reverse | CTCTGGGTTGGCACACACTT | Integrated DNA Technologies |
| ***IRF4*** | Forward | GAACGAGGAGAAGAGCATCTTCC | Integrated DNA Technologies |
| | Reverse | CGATGCCTTCTCGGAACTTTCC | Integrated DNA Technologies |
| ***nc209*** | Forward | GGTGGATCTGGTGTCCTCCT | Integrated DNA Technologies |
| | Reverse | ATTGGGTCTTTTCTCCGGCT | Integrated DNA Technologies |

**Table S3.** Top ten enriched pathways ranked by adjusted p-value in Cluster 9 samples.

| Pathway ID | Description | Gene Ratio | Fold Enrichment | P-adj. | Genes |
|---|---|---|---|---|---|
| **hsa04110** | Cell cycle | 71/183 | 2.30 | 7.3E-11 | CDKN2C/E2F2/TICRR/ESPL1/MCM2/PLK1/TRIP13/CDC20/AURKB/CCNA2/CDC45/CCNB1/CDK1/SMC1B/BUB1/TTK/CDCA5/CCNB2/ESCO2/PKMYT1/SGO1/CDC25C/CDC7/CDC25A/CDT1/MCM4/MCM3/BUB1B/E2F1/CDC6/KNL1/NDC80/ORC1/MAD2L1/MCM6/ORC6/PCNA/PTTG1/RBL1/DBF4B/MCM5/CHEK2/SMC1A/MTBP/CHEK1/FBXO5/CDK6/TFDP1/MCM7/ANAPC7/ANAPC1/SKP2/HDAC2/CCNB3/E2F3/RAD21/PPP2R5D/CDKN2A/ORC3/CDK2/YWHAG/CDK4/ORC5/PPP2R1B/YWHAE/HDAC8/RBL2/GSK3B/HDAC1/CDC23/ORC4 |
| **hsa03460** | Fanconi anemia pathway | 35/183 | 3.32 | 7.3E-11 | UBE2T/BRCA1/EME1/BRIP1/RMI2/FANCI/SLX4/FANCA/RMI1/RAD51/CENPS/CENPS-CORT/FANCB/RAD51C/FANCF/TOP3A/BLM/FAAP24/PALB2/RPA1/FANCE/USP1/RPA3/PMS2/FANCD2/FANCC/TELO2/MLH1/ERCC4/ATRIP/FANCM/REV3L/FAN1/TOP3B/FANCG |
| **hsa03030** | DNA replication | 26/183 | 3.70 | 1.2E-09 | MCM2/MCM4/MCM3/MCM6/PCNA/PRIM1/POLE2/POLA1/FEN1/PRIM2/RFC3/POLD1/POLD3/RFC5/DNA2/MCM5/RFC4/POLE/RFC2/POLD2/MCM7/RNASEH2A/RPA1/RPA3/LIG1/RNASEH1 |
| **hsa01240** | Biosynthesis of cofactors | 59/183 | 1.96 | 2.8E-06 | UGT2A1/UGT2A2/KYNU/AK7/HSD17B6/PANK1/GGH/NQO1/NME7/CTPS1/CAD/UMPS/PDXP/DHFR/COQ3/FLAD1/ADSL/PPOX/AK8/EARS2/KMO/MTHFD2/PSAT1/GCLM/ADSS2/UGDH/VKORC1L1/CPOX/PHOSPHO2/GSS/SHMT2/COQ5/COX15/DHODH/AFMID/UROS/NFS1/COQ7/ALDH1B1/DHFR2/PNPO/GGCX/SHMT1/MTHFD1/MPI/SPR/COQ2/COX10/ALAS1/DLD/NMNAT1/AK2/GMPPA/FECH/RFK/PANK3/NME6/PMM2/OXSM |
| **hsa03440** | Homologous recombination | 24/183 | 3.00 | 2.8E-06 | XRCC2/RAD54L/BRCA1/RAD54B/EME1/BRIP1/POLD1/POLD3/RAD51/RBBP8/RAD51C/RAD51D/TOP3A/BLM/POLD2/BRCC3/PALB2/TOPBP1/RPA1/RPA3/BARD1/RAD50/RAD52/TOP3B |
| **hsa03410** | Base excision repair | 25/183 | 2.91 | 2.8E-06 | NEIL3/PCNA/UNG/POLE2/FEN1/RFC3/POLD1/POLD3/RFC5/RFC4/POLE/RFC2/POLD2/PARG/LIG3/NEIL2/TDG/SMUG1/POLG2/LIG1/PARP4/MUTYH/ADPRS/PARP2/APTX |
| **hsa03430** | Mismatch repair | 16/183 | 3.57 | 1.3E-05 | EXO1/MSH2/PCNA/RFC3/POLD1/POLD3/RFC5/RFC4/MSH6/RFC2/POLD2/RPA1/RPA3/PMS2/LIG1/MLH1 |
| **hsa01200** | Carbon metabolism | 44/183 | 1.94 | 1.3E-04 | PRPS1L1/HKDC1/PFKP/ENO4/HAO1/CPS1/GLDC/ME3/GPT2/PSPH/HK2/SDSL/FH/SDHC/GOT2/PSAT1/PGP/PHGDH/ME2/SHMT2/ENO3/ACSS2/PCCB/PRPS1/PRPS2/RPE/SHMT1/MDH1/PDHA1/ACAT2/H6PD/DLD/ADH5/GOT1/HIBCH/RPIA/PDHB/SUCLA2/DLAT/PGAM1/MMUT/ECHS1/TKFC/MDH2 |
| **hsa04814** | Motor proteins | 64/183 | 1.69 | 2.1E-04 | TUBA3C/MYO18B/KIF7/KIFC3/TNNI1/MYH1/MYH14/DNAH2/MYH2/KIF14/KIF4A/KIF18B/DNAH14/KIF21B/MYH13/DNAH9/KIF20A/KIF2C/TUBB4A/MYO16/CENPE/MYO3A/KIF23/MYH8/KIFC1/DNAH5/KIF1A/MYH4/KIF15/KIF11/DYNLRB2/KIF24/DNAH7/DNAH3/DYNC2H1/KIF5C/DYNC1I1/KIF18A/TUBG1/TUBB/DNAH10/KIF22/KIF20B/KIFAP3/TPM2/BICDL1/TUBA1C/ACTA1/KIF5A/MYO19/DNAH6/DYNC2LI1/DNAI4/MYO1H/DYNC1I2/DYNLL2/MYL6B/DNAL4/DNAL1/DCTN1/KIF27/KIF2A/KIF3B/ACTR1A |
| **hsa04146** | Peroxisome | 33/183 | 2.04 | 5.5E-04 | DDO/PEX5L/HAO1/XDH/AMACR/DAO/SLC27A2/PECR/EPHX2/PXMP2/ABCD2/EHHADH/PEX11A/PEX19/DECR2/GNPAT/CROT/PMVK/AGPS/PRDX1/MPV17/SLC25A17/PEX11B/ACSL4/ABCD3/SOD1/FAR1/PEX26/HMGCL/ECI2/ACOT8/PEX13/PEX10 |

**Table S4.** Top ten enriched pathways ranked by adjusted p-value in Cluster 12 samples.

| Pathway ID | Description | Gene Ratio | Fold Enrichment | P-adj. | Genes |
|---|---|---|---|---|---|
| hsa04517 | IgSF CAM signaling | 37/607 | 1.91 | 0.017 | SLITRK4/SLITRK2/KCNA1/PAK5/MPDZ/JAM3/ROBO2/FGFR1/MPP3/NLGN1/PRKCG/MYH11/MAGI1/SCN8A/NTN1/PTPRS/LRRC4C/MPZ/PVR/ESAM/NFASC/PAK3/ROBO3/TUBA8/AKT3/TUBB6/GRIP2/PARD6G/CADM4/VAV3/CD48/SRGAP1/TUBA3D/MYLK3/LRRC4B/MYH10/DYNLL2 |
| hsa04012 | ErbB signaling pathway | 13/607 | 2.34 | 0.160 | NRG1/PAK5/PRKCG/SHC2/PAK3/SHC3/AKT3/SHC4/STAT5B/SHC1/ERBB2/CAMK2D/MTOR |
| hsa04550 | Signaling pathways regulating pluripotency of stem cells | 18/607 | 1.93 | 0.163 | FGFR2/NODAL/FZD8/FGFR1/MEIS1/ONECUT1/FZD2/BMP4/WNT3A/LIFR/AKT3/IGF1R/NANOG/NANOGP8/ESRRB/PCGF3/WNT2B/BMPR2 |
| hsa04114 | Oocyte meiosis | 17/607 | 1.90 | 0.185 | ADCY2/CPEB1/PGR/AR/IGF1R/SPDYC/SPDYE10/SPDYE13/SPDYE2/SPDYE6/CAMK2D/SPDYA/CPEB3/RPS6KA3/SPDYE1/PPP3CA/PPP3CB |
| hsa00760 | Nicotinate and nicotinamide metabolism | 7/607 | 2.85 | 0.218 | NT5E/SIRT4/PNP/ASPDH/NMNAT1/NT5C1B/SIRT5 |
| hsa01240 | Biosynthesis of cofactors | 18/607 | 1.81 | 0.218 | AK5/CMPK2/KMO/AK8/HPD/MTHFD2L/ASPDH/TPK1/LIPT1/COQ7/DHFR2/PPOX/GPHN/NMNAT1/DHODH/UGDH/GMPPA/COX15 |
| hsa04510 | Focal adhesion | 22/607 | 1.67 | 0.218 | LAMA1/PAK5/PRKCG/SHC2/MYL11/EMP2/PAK3/COL6A2/SHC3/AKT3/SHC4/IGF1R/TNXB/TLN2/THBS2/SHC1/VAV3/MYLK3/ITGA2/ERBB2/PIP5K1A/ROCK2 |
| hsa04390 | Hippo signaling pathway | 18/607 | 1.77 | 0.218 | FZD8/PPP2R2B/BMP2/FZD2/TCF7L1/GDF5/BMP4/WNT3A/TGFB2/GDF6/CTNNA3/PARD6G/BMP8B/DLG5/BMP8A/WNT2B/BMPR2/DLG3 |
| hsa04020 | Calcium signaling pathway | 26/607 | 1.58 | 0.218 | ADCY2/FGFR2/FGFR1/MCOLN3/RYR2/PRKCG/CACNA1A/FGF9/ADRB1/CHRNA7/MCOLN2/CACNA1E/CACNA1F/SLC8A3/NOS2/HTR4/GRPR/MYLK3/ERBB2/PLCD4/CAMK2D/ATP2A1/NFATC3/ARLN/PPP3CA/PPP3CB |
| hsa01521 | EGFR tyrosine kinase inhibitor resistance | 11/607 | 2.13 | 0.218 | FGFR2/NRG1/PRKCG/SHC2/SHC3/AKT3/SHC4/IGF1R/SHC1/ERBB2/MTOR |

#### Supplemental Figures

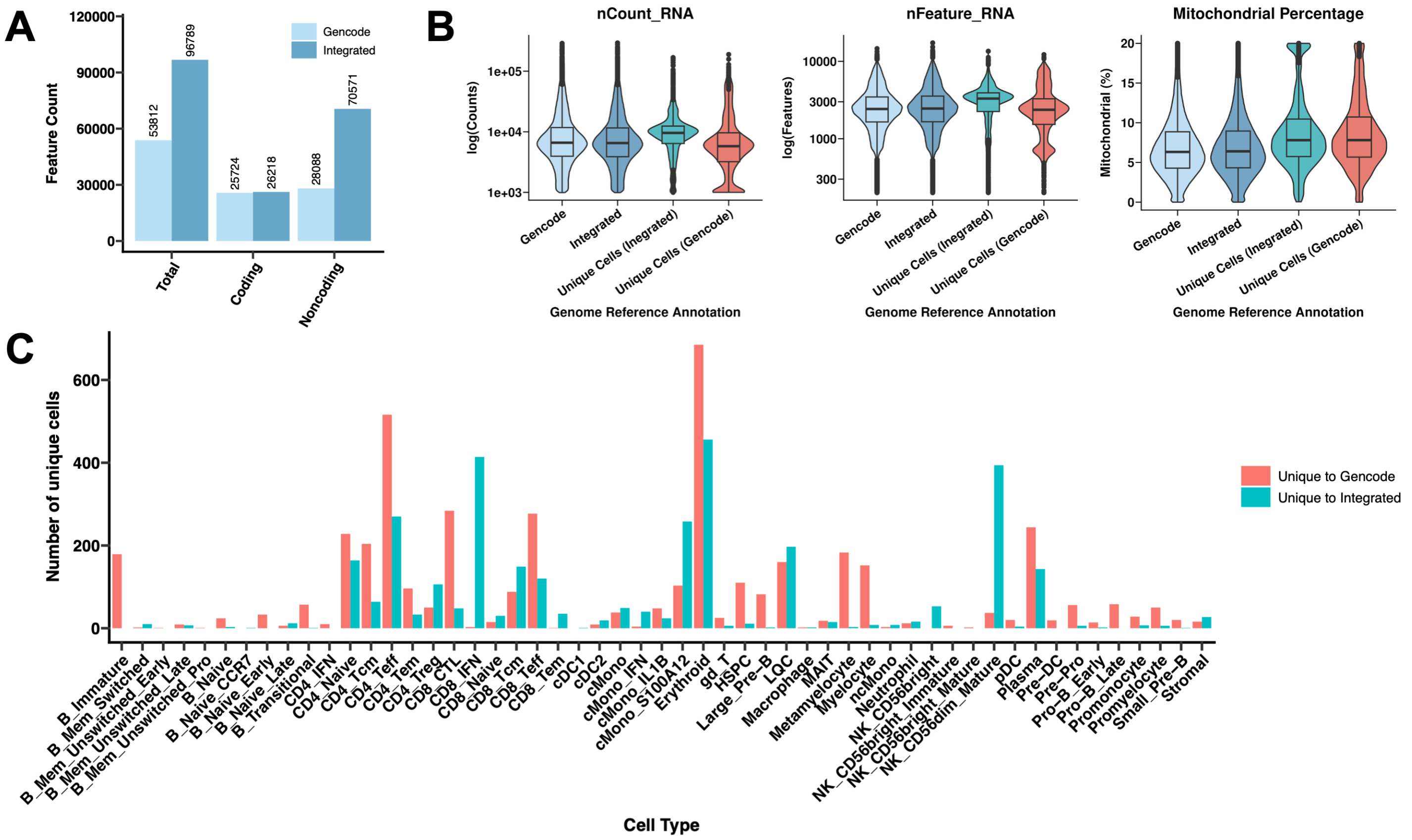


**Figure S1. (A)** Bar plot displaying the difference in the total number of unique coding and noncoding genes captured using the GENCODE v47 reference versus our integrated reference genome annotation. **(B)** Violin plots comparing standard quality control metrics for cells present in or unique to the benchmarking single-cell dataset after alignment using the GENCODE v47 reference versus our integrated reference genome annotation. **(C)** Bar plot representing the total number of each cell type present among mutually exclusive cells when aligning to the GENCODE v47 reference versus our integrated reference genome annotation.

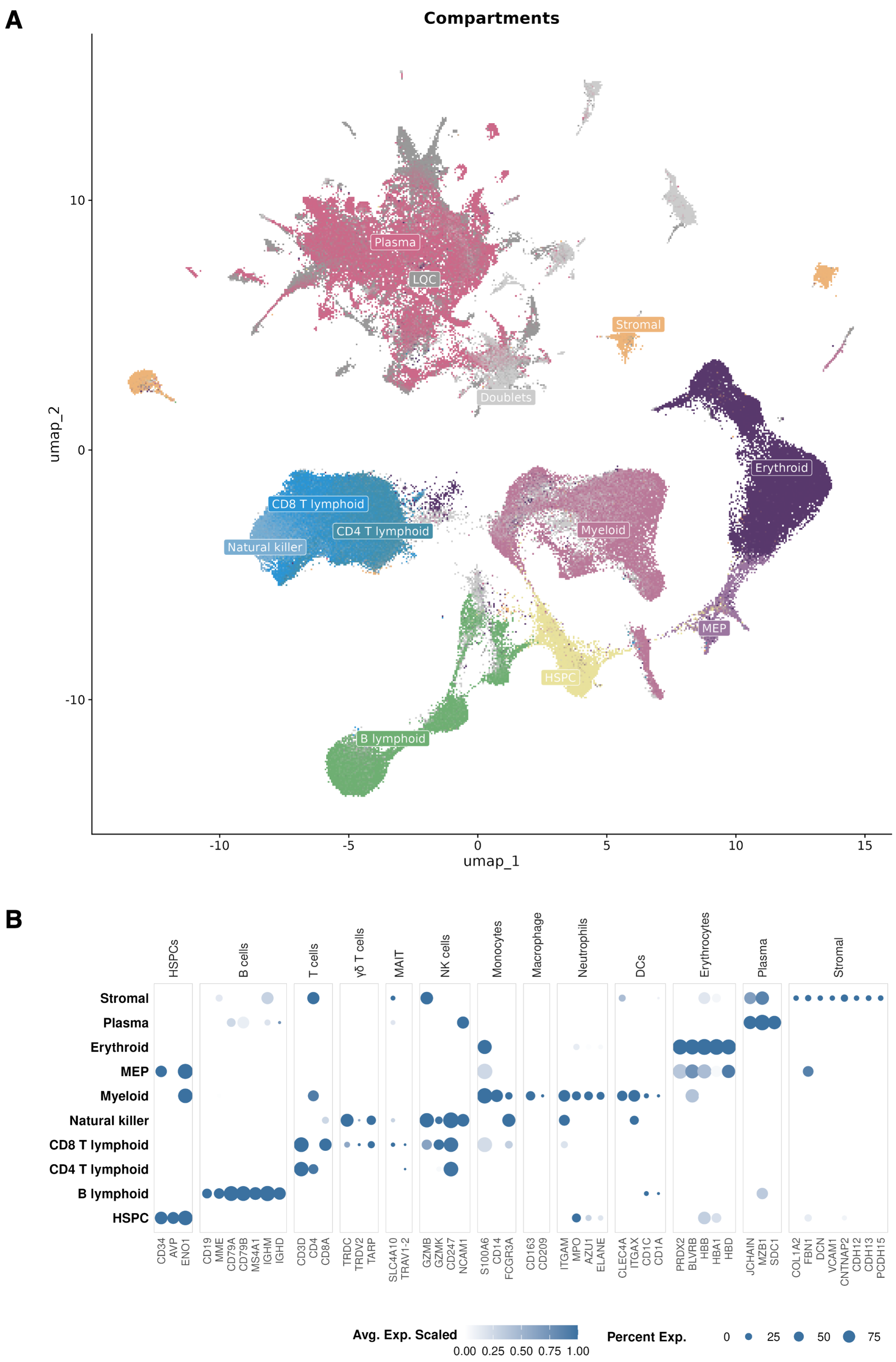


**Figure S2. (A)** UMAP embedding of 2,092,670 cells collected from CD138-negative sorted bone marrow aspirate samples. A total of 106 clusters were observed, spanning NK and T lymphoid, B lymphoid, erythroid, progenitor, stromal, and plasma cell lineages. **(B)** Dot plots displaying the average scaled gene expression for canonical lineage markers.

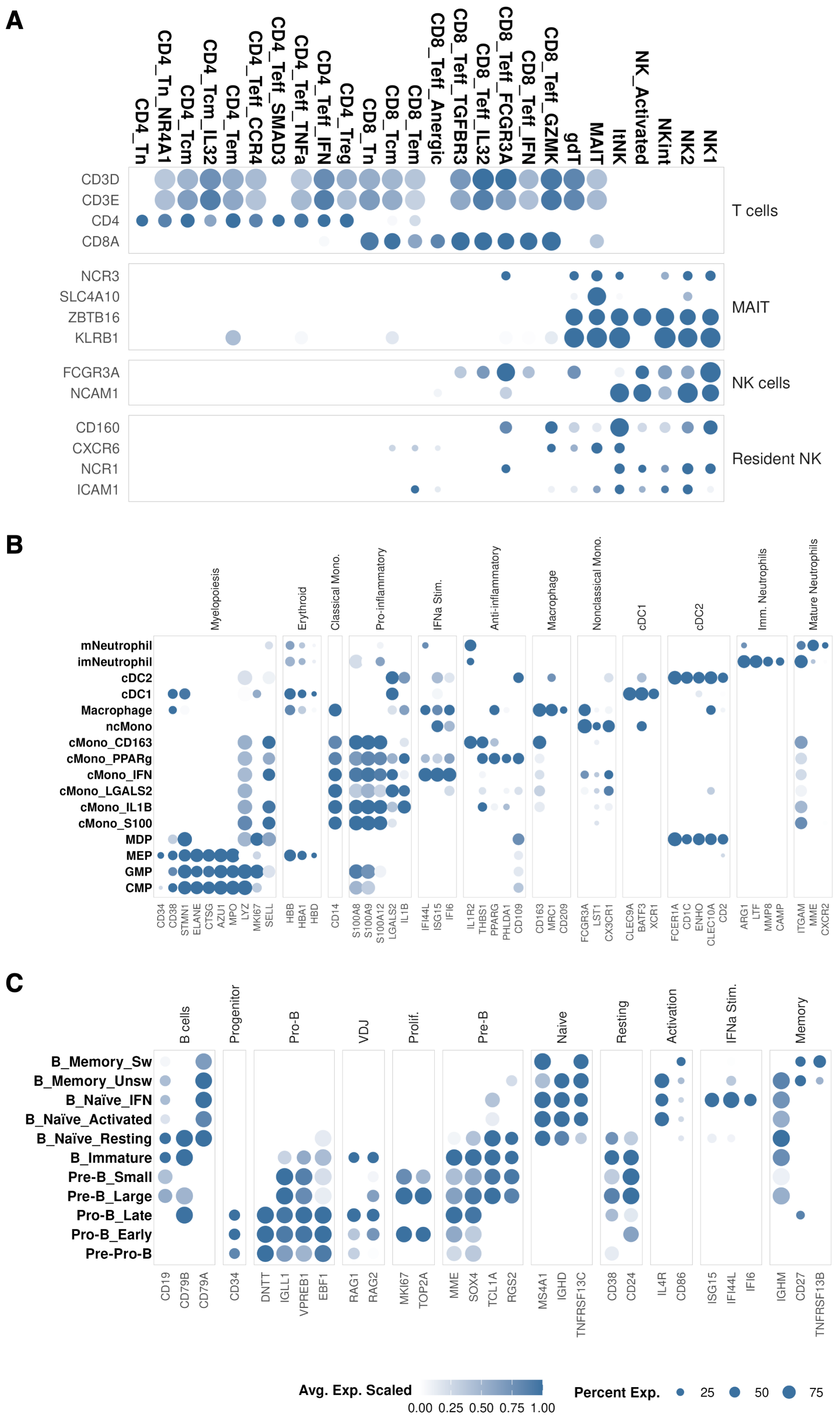


**Figure S3.** Dot plots displaying the average scaled gene expression of canonical markers for **(A)** NK and T lymphoid, **(B)** myeloid, and **(C)** B lymphoid subpopulations.

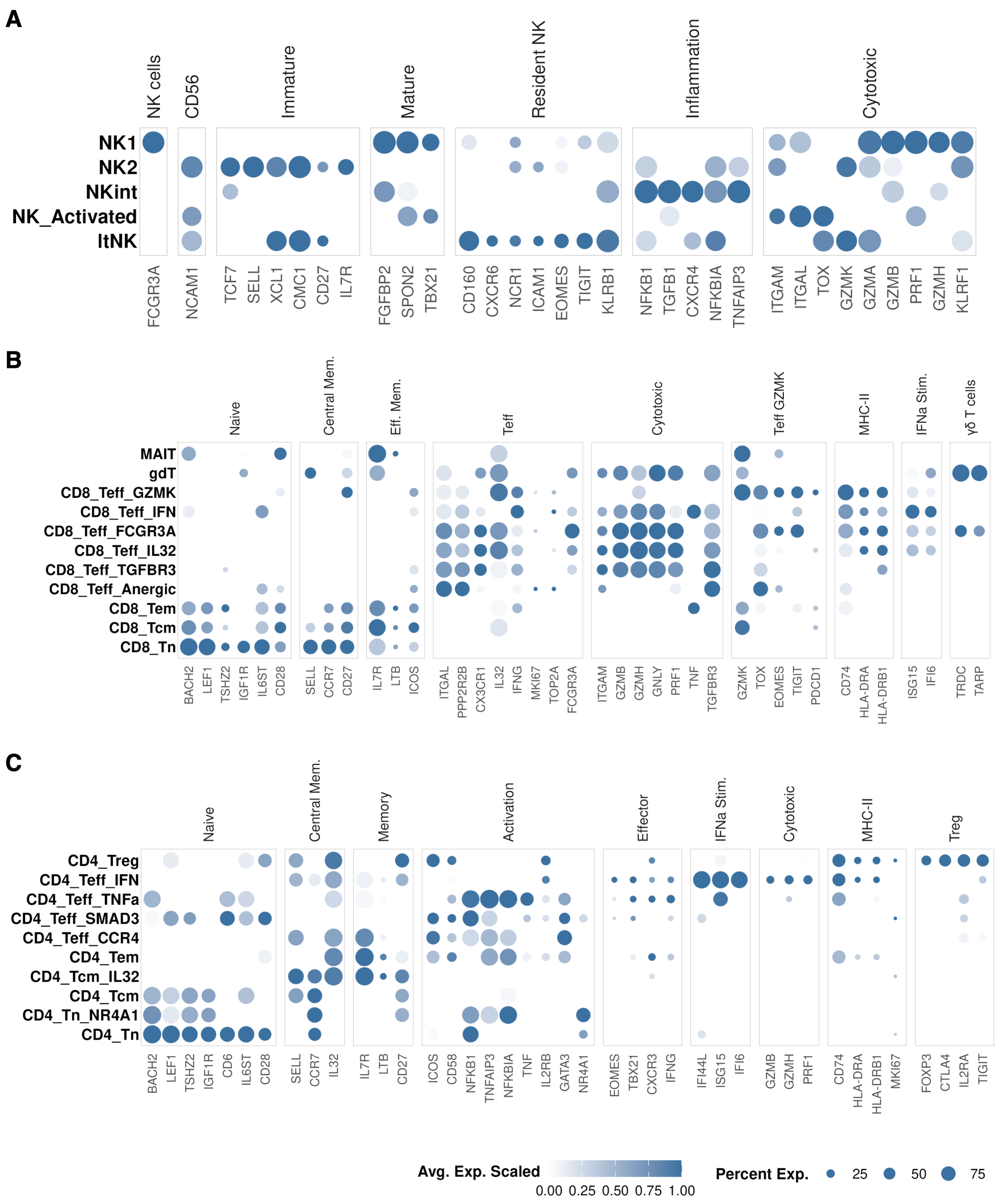


**Figure S4.** Dot plots displaying the average scaled gene expression of canonical markers for **(A)** NK, **(B)** CD8 T, and **(C)** CD4 T cell subpopulations.

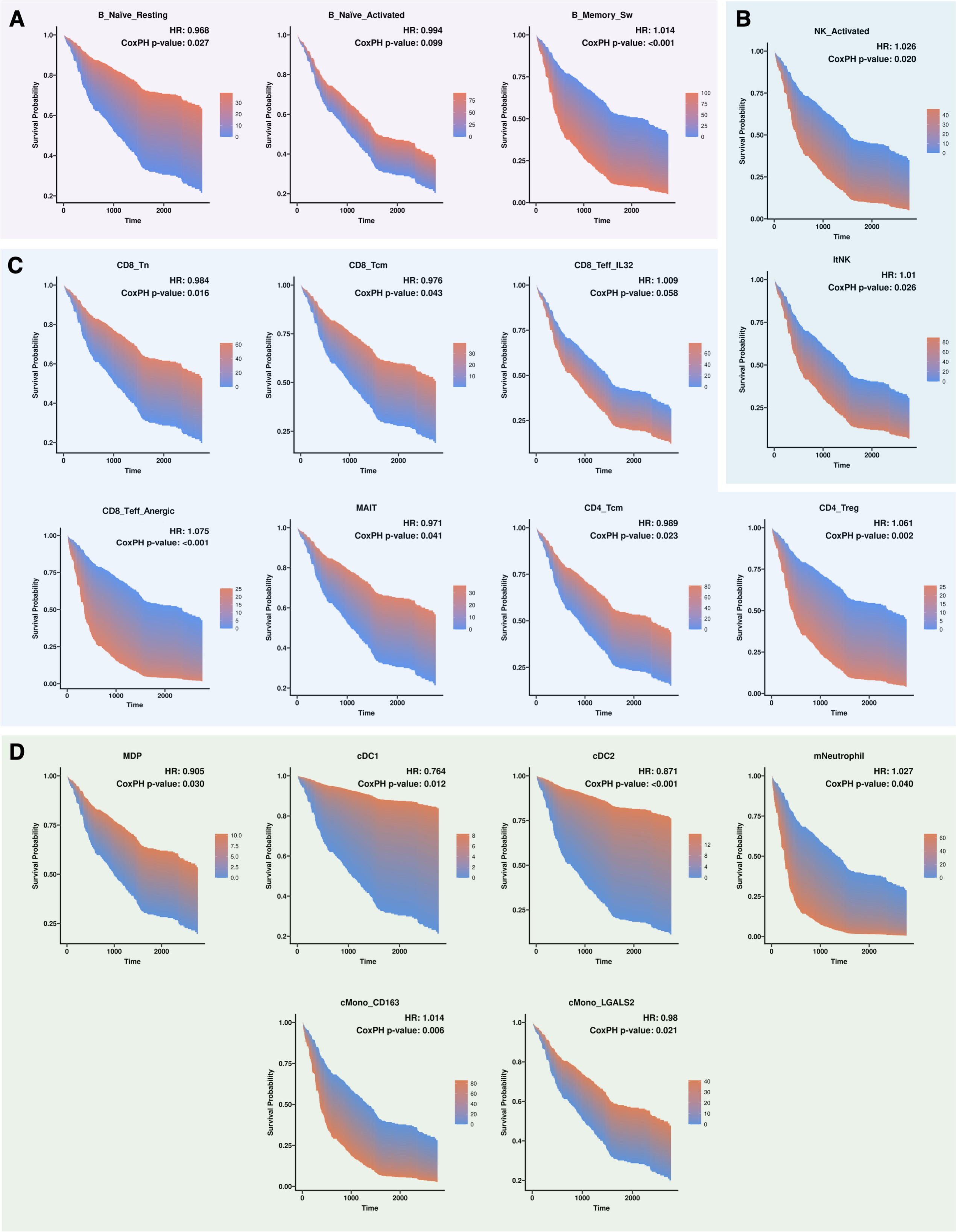


**Figure S5.** Kaplan-Meier curves for subpopulations exhibiting significant (P < 0.05) associations with progression-free survival across **(A)** B lymphoid, **(B)** NK, **(C)** T lymphoid, and **(D)** myeloid compartments.

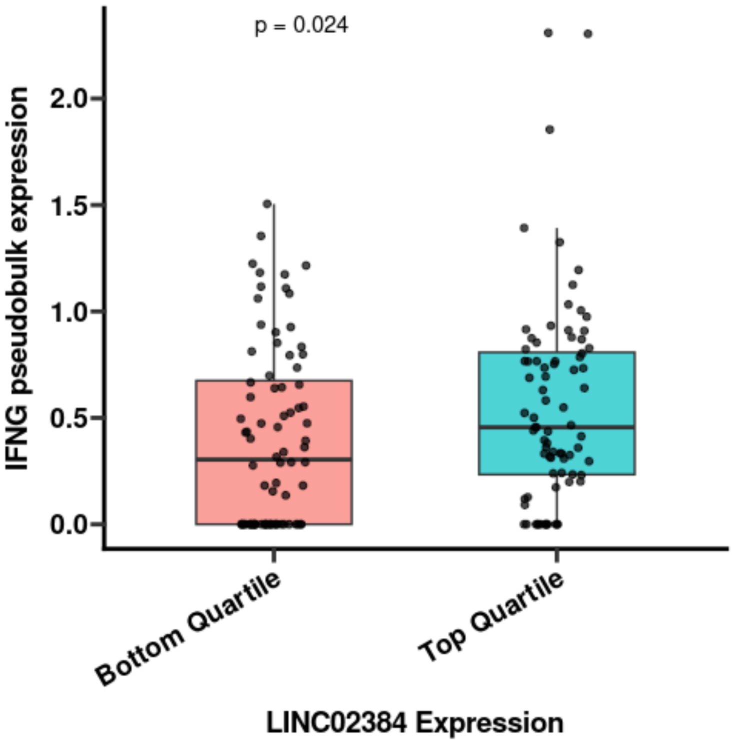


**Figure S6.** Box plots showing the difference in per-patient pseudobulk expression of interferon gamma (*IFNG*) between the top and bottom quartiles of samples expressing *LINC02384*.

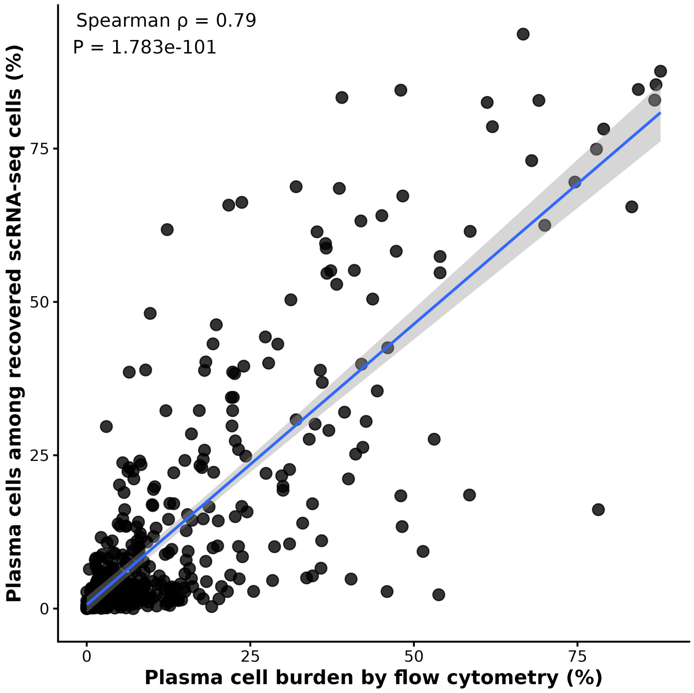


**Figure S7.** Correlation plot showing the relationship between plasma cell burden determined by flow cytometry and the proportion of plasma cells within the single-cell samples.

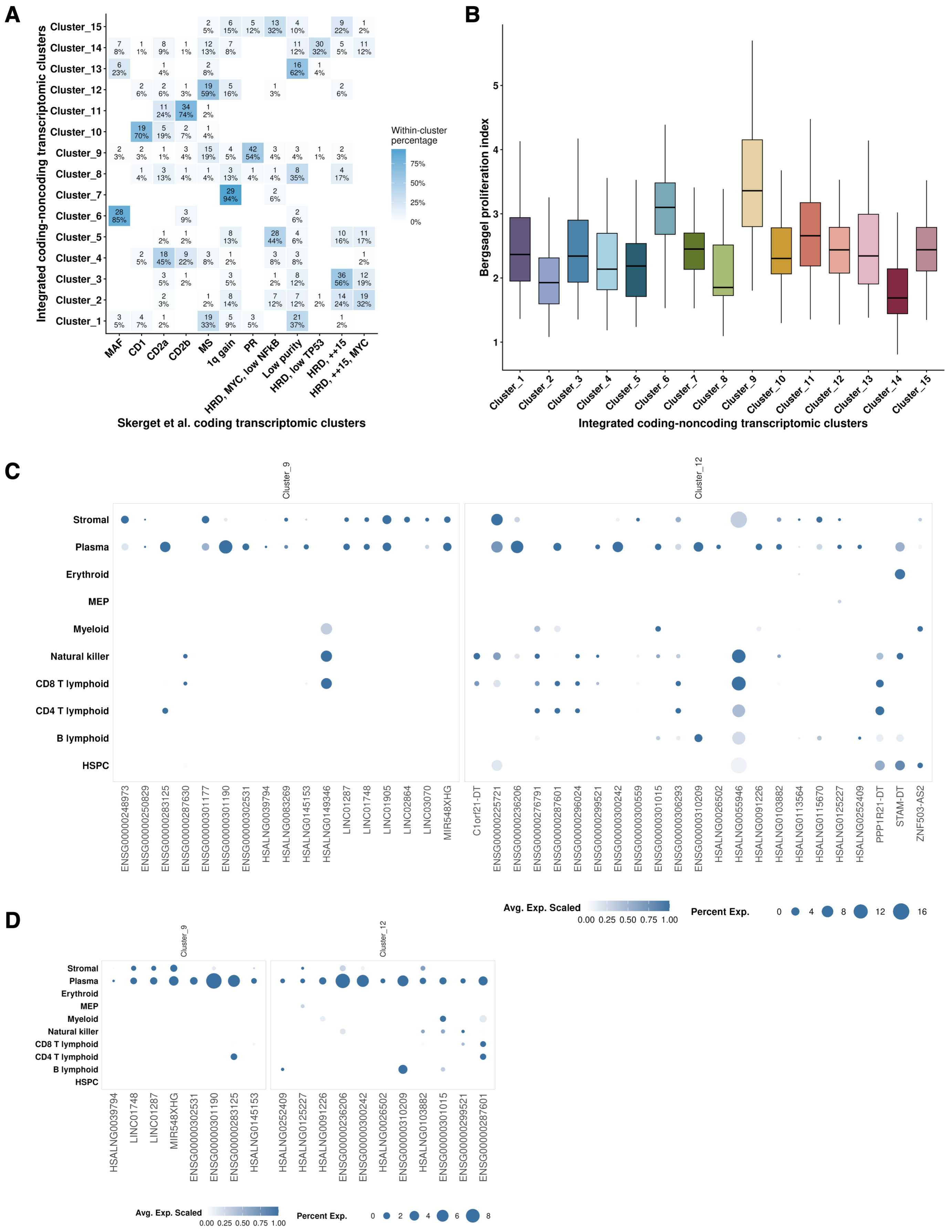


**Figure S8. (A)** Tile plot comparing cluster assignment using our integrated coding-noncoding genome annotation on the y-axis with cluster assignment reported by Skerget et al. using coding expression profiles for the same samples on the x-axis. [1] The clusters defined by Skerget et al. include: MAF family transcription factor expressing (MAF) characterized by t(14;16) patients, cyclin D expressing group 1 (CD1) characterized by t(11;14) patients, cyclin D expressing group 2a (CD2a) characterized by D-type cyclin IgH translocation patients, cyclin D expressing group 2b (CD2b) characterized by D-type cyclin targeting translocation patients, MMSET expressing (MS) characterized by t(4;14) patients, 1q gain characterized by patients with gain of chr1q, proliferation (PR) characterized by patients with a high proliferation index, and four high risk disease (HRD) clusters characterized by tetrasomy 15 (++15), structural events involving *MYC*, and low expression of *TP53*. **(B)** Box plot showing the average proliferation index for each derived patient cluster. **(C-D)** Dot plots examining the malignant versus non-malignant cell expression for **(C)** the candidate ncRNAs retaining concordant cluster specificity between the single-cell and bulk RNA-seq datasets and **(D)** the final 19 candidate ncRNAs that satisfied all quality-control criteria.

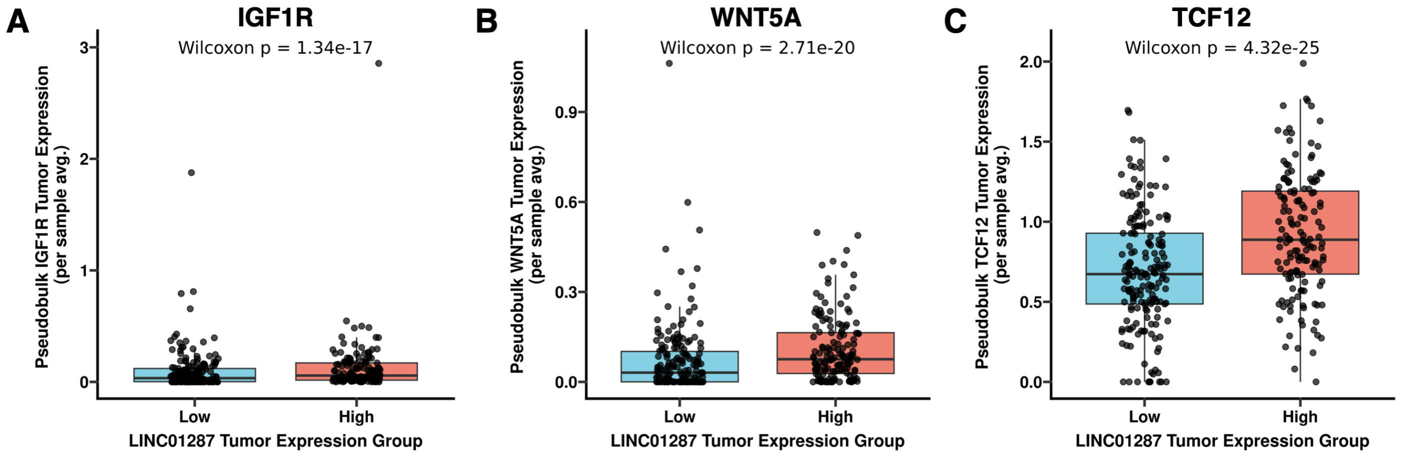


**Figure S9.** Box plots showing the increased expression of the previously reported mRNA targets of *LINC01287* **(A)** *IGF1R*, **(B)** *WNT5A*, and **(C)** *TCF12* in patients with high versus low *LINC01287* expression.[2–4]

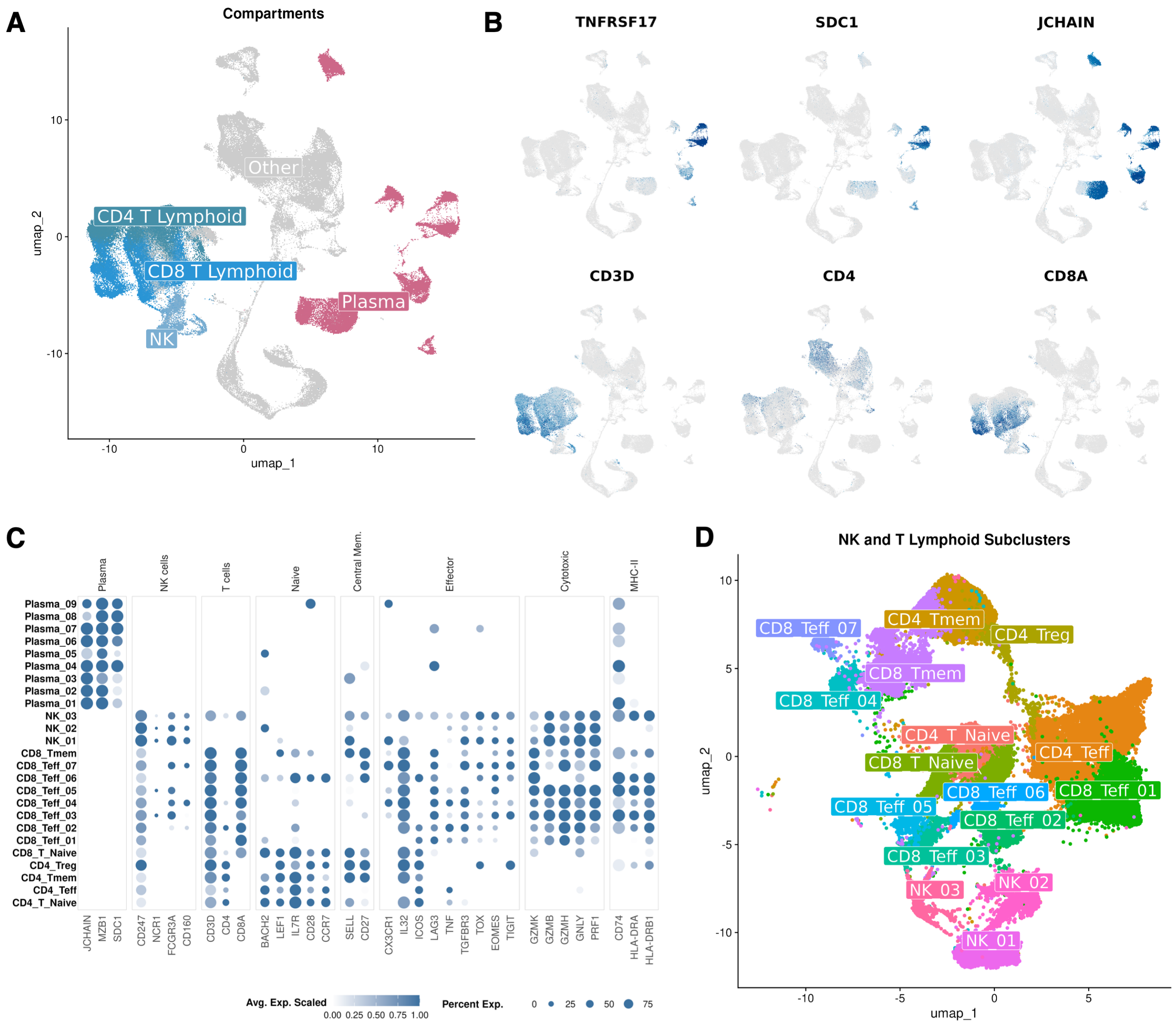


**Figure S10. (A)** UMAP embedding of 133,223 cells generated from the Dhodapkar et al. dataset (GSE210079) of bone marrow aspirate samples taken before and after CAR-T therapy. Cells are colored based on their lineage compartment. **(B)** Feature plots projecting the scaled expression of key plasma and T cell lineage markers onto UMAP embeddings. **(C)** Dot plot displaying the average scaled gene expression for canonical markers of plasma, NK, and T cell lineages. **(D)** UMAP embedding of the 46,720 cells from NK, CD4 T, and CD8 T lymphoid compartments, colored by subcluster.

### Supplemental Methods

#### Method S1. Development of an integrated coding-noncoding reference genome annotation

##### 1.1 Pipeline for systematically merging reference genome annotations

A five-step computational pipeline using Bash and Python was developed to integrate standard and noncoding reference genome annotation files into unified coding-noncoding reference annotations. We applied this pipeline to systematically integrate the noncoding reference annotation LncBook v2.1 with GENCODE v47,[5,6] expanding the noncoding coverage of the genome annotation while strictly preserving coding genome annotation.

First, the original GENCODE and LncBook GTF files were standardized using GffRead (v0.12.8) to resolve malformed features, consolidate compatible transcript and exon records, retain exon-level attributes, generate exon features for gene-only records, and remove duplicate sequences.[7] The resulting GTF files were validated to ensure checks for valid features, correct start and end positions, strand consistency (removing records without annotated strands), score validity, frame correctness, and proper attribute formatting. To prioritize ncRNA annotations already present in the GENCODE reference, GENCODE transcripts annotated as scaRNA, scRNA, snoRNA, snRNA, sRNA, lncRNA, miRNA, or miscellaneous RNA were separated from the remaining coding GENCODE annotation. The noncoding LncBook transcripts were compared with these GENCODE ncRNA transcripts using GffCompare (v0.12.9),[7] and only LncBook transcripts assigned class codes "i," "p," or "u," representing intronic, potentially intergenic, or otherwise non-overlapping transcript models, were retained. These nonredundant LncBook transcripts were combined with the original GENCODE ncRNA annotations to generate a noncoding genome annotation, which was then compared with the GENCODE coding genome annotation using GffCompare in Step 2 to classify each ncRNA transcript according to its genomic relationship with coding transcripts. Noncoding transcripts assigned class codes "i," "p," or "u" were retained without modification. Transcripts assigned class codes "s" or "x," indicating antisense overlap, and class codes "o" or "y," indicating sense overlap, were retained for overlap resolution in Steps 3-4. Noncoding transcripts assigned other class codes, representing ncRNAs identical to, contained within, or otherwise structurally overlapping coding transcripts, were subsequently removed.

For the sense- and antisense-overlapping ncRNAs, regions shared with coding features were removed using BEDTools (v2.31.1) subtract in Step 3.[8] Sense-strand overlaps were resolved using strand-specific subtraction, whereas antisense overlaps were resolved by subtracting coding features on the opposite strand. The resulting GTF files were cleaned and validated using GffRead, and duplicate GTF records introduced by feature splitting were removed. In Step 4, an additional 250-bp segment was removed from each trimmed boundary to reduce ambiguous read assignment near coding-noncoding junctions. Buffered transcript intervals that had a remaining coordinate span of 100 bp or less were discarded. These thresholds were selected based on the average short read sequencing lengths typically used to profile RNA-sequencing libraries. The terminal exon coordinates for trimmed noncoding transcripts were constrained to reflect their updated transcript boundaries.

##### 1.2 Generation of final integrated coding-noncoding human reference genome annotation

In Step 5, the unmodified non-overlapping ncRNA transcripts and the trimmed and buffered sense- and antisense-overlapping ncRNA transcripts were concatenated and processed with GffRead to consolidate compatible records, remove duplicate features, and validate the resulting transcript structures. This finalized ncRNA annotation was then concatenated with the GENCODE coding annotation, followed by final validation using GffRead with forced exon generation to ensure that each retained transcript was represented by valid transcript and exon features. Selected biotypes that were not relevant to the intended transcriptomic analysis or that could introduce low-confidence or redundant features were removed. These included artifacts, transcripts subject to non-stop decay, transcripts annotated as "to be experimentally confirmed," mitochondrial ribosomal and transfer RNAs, vault RNAs, ribosomal RNAs, ribozymes, T-cell receptor gene segments, selected immunoglobulin pseudogenes, processed and unitary pseudogenes, and pseudogene classes. The resulting GTF constituted the final integrated coding-noncoding human reference annotation.

Gene and transcript composition of the integrated reference annotation was summarized directly from the final GTF. Noncoding genes were labeled according to whether they originated from the unmodified, sense-overlapping, or antisense-overlapping annotation sets. When a gene was represented in more than one noncoding category, it was assigned using the priority order sense, antisense, and non-overlapping.

##### 1.3 Alignment of RNA-sequencing data to the integrated human reference genome annotation

A Cell Ranger-compatible transcriptome reference was generated from the GRCh38.p14 genomic FASTA and the final integrated GTF using the 10x Genomics Cell Ranger v8.0.1 mkref command.[9] The resulting reference was used to align all 10x Genomics single-cell RNA-sequencing raw FASTQ files using Cell Ranger count. A separate STAR genome index was generated from the same GRCh38.p14 assembly and integrated GTF for alignment of the 942 paired-end CD138-positive bulk RNA-sequencing FASTQ files from the MMRF CoMMpass study using STAR (v2.7.11b).[10] STAR was run with the integrated genome index using the parameters --quantMode TranscriptomeSAM GeneCounts.

#### Method S2. Benchmarking of the integrated coding-noncoding reference genome annotation

##### 2.1 Generating the benchmarking datasets

A benchmarking dataset was generated by randomly selecting 12 samples from each of the four batches of single-cell RNA-sequencing data generated from bone marrow aspirates of patients enrolled in the MMRF CoMMpass study. To evaluate the performance of the integrated coding-noncoding reference annotation, the raw single-cell RNA-sequencing data from the selected 48 samples were independently aligned to three human reference annotations: GENCODE v47, a direct concatenation of GENCODE v47 and LncBook v2.1 without resolution of overlapping loci, and the integrated GENCODE v47-LncBook v2.1 annotation. Reference transcriptomes were generated using 10x Genomics Cell Ranger v8.0.1 with the same GRCh38 genome assembly.[9] Raw sequencing data from each library were processed using cellranger count, and the resulting raw feature-barcode matrices were imported into Scanpy (v1.9.3),[11] filtering for cells with >1,000 total unique molecular identifier (UMI) counts, at least 200 detected genes, and less than 20% mitochondrial RNA. Filtered count matrices were then imported into Seurat (v5.4.0).[12] Genes were retained for downstream analysis if they had more than 10 total UMIs, were detected in more than 10 cells, and were expressed in more than 20% of the 48 samples.

##### 2.2 Benchmarking metrics

To determine whether the integrated annotation altered dimensionality reduction, clustering, and annotation, the GENCODE- and integrated-aligned datasets were processed independently using identical parameters. The percentage of ambiguous reads per sample was calculated using the Cell Ranger "Reads Mapped Confidently to Transcriptome" metric. Cell recovery was compared by matching cell barcodes within each sample and quantifying cells shared between, or uniquely retained by, the GENCODE- and integrated-aligned datasets. Counts were normalized using Seurat's LogNormalize function, and the 3,000 most highly variable genes were identified using the variance-stabilizing transformation method, regressing immunoglobulin and ribosomal genes. Principal component analysis (PCA) was then performed using the scaled variable gene expression. Batch-correction was performed using Harmony (v.2.0.5),[13] correcting for batch by study site. Shared nearest-neighbor graphs were constructed using the first 50 Harmony dimensions, followed by graph-based clustering at a resolution of 1.0 and uniform manifold approximation and projection using the same dimensions. Several samples contained murine cell contaminants from the original study;[14] clusters identified as mouse-derived on the basis of species-specific marker expression were removed.

Major cellular compartments were annotated using canonical lineage markers identified with Seurat's FindAllMarkers function. Immune compartments were subsequently subclustered to identify immune subpopulations. B cells were subclustered using 20 Harmony dimensions at a resolution of 1.0, whereas myeloid and NK/T cells were subclustered using 40 and 30 Harmony dimensions, respectively, at a resolution of 0.75. Subclusters were manually annotated using established lineage, differentiation, and functional markers. Across all compartments, differential expression between the annotated subpopulations was performed using Seurat's FindAllMarkers function and the Wilcoxon rank-sum test with Bonferroni correction.

#### Method 3. Single-cell RNA-sequencing dataset processing and analysis

##### 3.1 Pre-processing and quality control

Raw count matrices generated by Cell Ranger were imported into Scanpy from the raw_feature_bc_matrix output for each sample. Cells were retained if they contained >1,000 UMI counts, > 200 unique features, and <20% mitochondrial transcript abundance. The resulting quality-controlled matrices were concatenated across samples and converted to a Seurat v5 object using BPCells (v0.3.1) to support memory-efficient analysis.

Genes in the final count matrix were retained if they had more than 10 total counts, were detected in more than 10 cells, and were expressed in more than 20% of samples. Counts were normalized using Seurat's LogNormalize method. The 3,000 most variable features were identified, after which immunoglobulin and ribosomal protein genes were excluded from the variable-feature set to reduce clustering driven by immunoglobulin abundance or ribosomal expression.

Due to the size of the combined dataset, a representative sketch containing 10% of all cells was generated using Seurat's SketchData function. The variable features were scaled within the sketch assay, and principal component analysis was performed using 50 principal components.

##### 3.2 Clustering and coarse cell type annotation

Batch correction was performed in the sketched dataset using Harmony. Harmony integration was performed over 50 principal components using the combined sequencing-site and batch variable. To promote convergence in this large dataset, Harmony was run with 100 k-means initializations, a maximum of 5,000 k-means initialization iterations, and a maximum of 50 Harmony iterations, with early stopping disabled.

A shared nearest-neighbor graph was constructed based on the first 30 Harmony dimensions and a resolution of 1.75, followed by graph-based clustering using Seurat, and a two-dimensional uniform manifold approximation and projection embedding was generated for visualization. Cluster assignments and the fitted UMAP model were projected from the sketch to the complete dataset using ProjectData. To remove the murine cells present in several samples from the original study,[14] murine-derived clusters were identified by their expression of epithelial markers, separation from human hematopoietic populations, and restricted representation across samples and were subsequently removed before final clustering.

Following removal of mouse-cell clusters, the sketch was reclustered using the first 50 Harmony dimensions and a resolution of 1.0. A shared nearest-neighbor graph was constructed, graph-based clustering was performed using Seurat, and a two-dimensional uniform manifold approximation and projection embedding was generated for visualization. Cluster assignments and the fitted UMAP model were projected from the sketch to the complete dataset using ProjectData. Coarse cell identities were assigned manually using cluster-enriched genes determined using Seurat's FindAllMarkers, canonical lineage markers, and previously defined cell labels from the MMRF Immune Atlas.[14] Clusters comprised of putative doublets and low-quality cell populations were identified based on mixed-lineage marker expression, elevated or atypical RNA content, and cluster-level quality-control metrics and were removed from the final annotated dataset.

##### 3.3 Immune compartment analysis

B lymphoid, myeloid, NK and T lymphoid compartments were analyzed separately to resolve distinct immune subpopulations. After subsetting the cells within each compartment, the 3,000 most variable features were identified, with immunoglobulin genes excluded from the variable-feature set. Expression values were scaled, and principal component analysis was performed using 50 principal components. Batch correction was repeated within each compartment using Harmony and the combined sequencing-site and batch variable. The B lymphoid and myeloid compartments were reclustered using 40 Harmony dimensions at a resolution of 0.6, while the NK and T lymphoid compartment was reclustered using 40 Harmony dimensions at a resolution of 1.0. UMAP embeddings were generated from the corresponding Harmony dimensions. Subclusters were annotated manually using canonical lineage, differentiation, activation, and cell-state markers. Subclusters exhibiting markers from multiple distinct lineages were designated as putative doublets, whereas clusters with low transcript complexity or nonspecific stress-associated expression were designated as low-quality cells. Both groups were excluded from downstream analyses.

Following annotation, differentially expressed genes among immune subtypes were identified using FindAllMarkers with significant genes defined as having a $\log_2$fold-change less than -1 or greater than 2, detection in greater than 10% of cells, and an adjusted P-value less than 0.05 using a Wilcoxon rank sum test with Bonferroni correction. Differentially expressed noncoding RNAs were identified by intersecting subtype marker lists with the integrated noncoding RNA annotation.

Associations between baseline immune-subpopulation abundance and progression-free survival were evaluated at the patient level. For each sample, the abundance of a subtype was calculated as its percentage of cells within the corresponding immune compartment. Cox proportional-hazards models were fitted with progression-free survival time and censoring status as the outcome, subtype percentage as a continuous predictor, and sequencing site/batch as a covariate. Where shown, high- and low-abundance groups were defined for Kaplan-Meier visualization using an optimal cutpoint threshold calculated from the survMisc (v.0.5.6) cutp function.

##### 3.4 *In silico* functional analysis of *ENSG00000310209*

To investigate the potential functional role of *ENSG00000310209*, plasma-cell counts were aggregated by sample to generate per-sample pseudobulk expression profiles. Samples were classified as "ENSG00000310209-high" or "ENSG00000310209-low" according to median *ENSG00000310209* expression unless otherwise specified. Differential expression of selected genes between groups was evaluated using Wilcoxon rank-sum tests with Bonferroni correction.

Plasma-cell gene co-expression networks were constructed using hdWGCNA (v0.4.11).[15] The input feature set comprised the 5,000 most variable genes together with the 19 candidate noncoding RNAs. Metacells were constructed within samples using 25 nearest neighbors and a maximum shared-neighbor value of 10. A signed co-expression network was generated using a soft-power threshold of 8. Following hdWGCNA, genes assigned to each co-expression module were converted to Entrez identifiers and tested for Reactome pathway over-representation using ReactomePA (v1.54.0).[16] Pathways associated with infectious or antimicrobial responses were excluded to prioritize biologically relevant signaling programs. Intramodular connectivity was quantified using module eigengene-based connectivity (kME), and the ten most highly connected genes in each module were designated as hub genes. Module networks were then visualized to examine relationships among hub genes and identify highly connected coding and noncoding transcripts.

To infer the activity of Wnt and IRF4 signaling pathways, pathway gene set enrichment from MSigDB via msigdbr (v26.1.0) was used.[17,18] Canonical Wnt signaling activity was evaluated using the GOBP_CANONICAL_WNT_SIGNALING_PATHWAY genes present in the single-cell dataset to calculate a Seurat module score, which was averaged by sample and compared between the ENSG00000310209 expression groups using a Wilcoxon rank-sum test with Benjamini-Hochberg correction. IRF4 activity was assessed by measuring the per-gene pseudobulk expression of genes present in the SHAFFER_IRF4_MULTIPLE_MYELOMA_PROGRAM gene set generated by Schaffer *et al*.[19] and comparing between the ENSG00000310209 expression groups using Wilcoxon rank-sum tests with Benjamini-Hochberg correction.

Outgoing intercellular signaling associated with nc209-high tumor cells was identified by differential expression using Seurat's FindMarkers contrasting plasma cells from samples in the upper quartile of *ENSG00000310209* expression and the remaining samples, followed by filtering to identify ligands using the CellChatDB interaction database (v2.2.0).[20] Significance was evaluated using Wilcoxon rank-sum tests with Benjamini-Hochberg correction.

To evaluate potential effects on the immune microenvironment, differential expression within the CD8+ T cell compartment was performed using Seurat's FindMarkers between ENSG00000310209 expression groups using Wilcoxon rank-sum tests with Benjamini-Hochberg correction. Markers of CD8+ T cell function and anergy were examined using Seurat's DotPlot function. CD8+ T cell differentiation trajectories were reconstructed with Monocle 3 (v.1.4.26).[21] Cells were clustered at a resolution of 5.0e-6, a principal graph was learned without partitioning or closed loops using 500 graph centers, and pseudotime was rooted in the naïve CD8+ T cell cluster. The expression of the immune-checkpoint genes *CD274* (PD-L1) and *PDCD1LG2* (PD-L2) was evaluated between ENSG00000310209 expression groups in malignant plasma cells using a Wilcoxon test.

##### 3.5 Differential abundance of immune subpopulations

The differential abundance of immune subpopulations based on the high versus low expression of each of the 19 candidate ncRNAs was calculated using Dirichlet regression with the DirichletReg R package (v0.7-2). Analyses were restricted to baseline samples. Plasma-cell counts were aggregated by sample, and patients were classified as high or low for each candidate noncoding RNA using the median pseudobulk expression, wherein expression above the cohort median was considered high and expression at or below the median was considered low. For each immune compartment, a sample-by-subtype count matrix was constructed. Zero counts were replaced with 0.5, and counts were divided by the total number of cells in each sample to produce compositional proportions. The most abundant subtype across samples was selected as the reference category. Differential abundance between ncRNA-high and ncRNA-low groups was evaluated using the DirichReg function, testing the association between abundance and expression group (i.e., immune-subtype composition~expression group). Subtype-specific Wald tests were then used to estimate the effect of high versus low expression relative to the most prevalent reference subtype and adjusted using the Benjamini-

Hochberg method. Effect estimates were reported on the log-odds scale, with corresponding odds ratios calculated by exponentiation.

##### 3.6 In silico validation using Dhodapkar *et al.* dataset (GSE210079)

Raw single-cell RNA-sequencing data originally reported by Dhodapkar *et al.* (GSE210079)[22] were reprocessed to evaluate whether the tumor-intrinsic and immune associations of nc209 were reproduced in an independent multiple myeloma cohort. The dataset comprised sequencing data from 23 bone marrow aspirates collected from 11 patients before treatment (N = 6), at day 28 (N = 6), or at least three months after treatment (N = 11). Publicly available BAM files were converted to FASTQ files using the bamtofastq function in Cell Ranger v8.0.1. The BAM file from patient 8 contained insufficient data to convert to FASTQ and was subsequently excluded from the dataset. The resulting FASTQ files were then processed using Cell Ranger against the integrated coding-noncoding reference. Gene-expression matrices were imported from the Cell Ranger filtered feature-barcode matrices and merged in Seurat.

Consistent with our internal dataset processing, cells were retained if they had more than 1,000 total counts, more than 200 detected genes, and less than 20% mitochondrial transcript abundance. Genes were retained if they had more than 10 total counts, were detected in more than 10 cells, and were expressed in more than 20% of libraries. Counts were normalized using LogNormalize with a scale factor of 10,000. The 3,000 most variable features were identified using the variance-stabilizing transformation method, with immunoglobulin and ribosomal protein genes excluded from the variable-feature set. Expression values were scaled, and principal component analysis was performed using 50 principal components.

Graph-based clustering was performed using Seurat with the first 40 principal components at a resolution of 0.8, generating 42 clusters, which were manually assigned to plasma, natural killer and T lymphoid, or “other” compartments based on canonical lineage markers.

Cells from the natural killer and T lymphoid compartment were subsequently reclustered to resolve CD8+ T cell subpopulations. Within this compartment, 3,000 variable features were identified, immunoglobulin and ribosomal genes were excluded, and principal component analysis was repeated using 50 components. A shared nearest-neighbor graph was generated using the first 30 principal components, followed by clustering at a resolution of 0.8 and manual annotation using canonical markers of CD4+ and CD8+ T cells, natural-killer cells, naïve and memory T cells, and cytotoxic and effector T cells.

Plasma-cell analyses were restricted to samples containing more than 10 annotated plasma cells (N = 13). Plasma-cell counts were aggregated by sample using Seurat’s PseudobulkExpression function. Samples were classified as nc209-high when their scaled pseudobulk *ENSG00000310209* expression was above the cohort median and as nc209-low when below the median.

Pseudobulk nc209 expression was compared between samples from patients with and without the t(4;14) translocation using a Wilcoxon rank-sum test. Differences in nc209 expression among the three clinical treatment response groups were evaluated using a Kruskal-Wallis test.

To validate Wnt and CCL5 signaling in this independent cohort, canonical Wnt pathway activity was quantified using the MSigDB GOBP_CANONICAL_WNT_SIGNALING_PATHWAY gene set to calculate a Seurat module score for each cell. The average canonical Wnt score per sample and pseudobulk *CCL5* expression were compared between nc209-high and nc209-low samples using Wilcoxon rank-sum tests with Benjamini-Hochberg correction.

Lastly, to investigate potential alterations to T cell transcriptional states associated with nc209 expression and treatment response, CD8+ T effector clusters were combined and compared between responder and non-responder samples. The differential expression of mRNAs was evaluated using Seurat's Wilcoxon rank-sum test with a minimum detection fraction of 5%.

#### Method S4. Bulk RNA-sequencing dataset processing and analysis

##### 4.1 Pre-processing and quality control

Gene-level counts for the 942 CD138-positive bulk RNA-sequencing samples from the MMRF CoMMpass study were obtained from the STAR ReadsPerGene.out.tab files using the unstranded count column and combined across samples into a single gene-by-sample count matrix. Sample identifiers were matched to clinical metadata, and all downstream analyses were restricted to samples collected at baseline (N = 776).

Genes were retained in the final count matrix if they had at least one read in 39 or more samples. Genes with total counts below the 1st percentile or above the 99th percentile of the gene-level count distribution were removed. Raw counts were imported into DESeq2 (v.1.50.2),[23] normalized using its median-of-ratios method for differential expression and transformed using the variance-stabilizing transformation with blind dispersion estimation for downstream clustering and survival-association analyses.

##### 4.2 Consensus clustering of patients based on tumor expression profiles

Patients were grouped according to their coding-noncoding tumor expression profiles using consensus clustering.[1] The top 10% most variable genes across baseline samples were selected from the variance-stabilized transformation expression matrix, and consensus partitioning around medoids clustering was performed using ConsensusClusterPlus (v.1.74.0).[24] Candidate solutions ranging from 2 to 30 clusters were evaluated using 100 resampling iterations, with 80% of patients included in each iteration, all selected genes retained, and Pearson correlation used as the distance metric. A k value of 15 was selected for downstream analysis, and the resulting patient groups were designated Cluster 1 through Cluster 15. A k value of 15 was selected based on the relative change in area under the cumulative distribution function (CDF) curve, while maximizing cluster resolution and ensuring that the smallest cluster contained at least 25 patients.

Associations between transcriptomic cluster membership and progression-free survival were assessed using Cox proportional-hazards regression. Each patient cluster was compared with all remaining patients, with adjustment for autologous stem-cell transplantation, race, and sex. Kaplan-Meier curves and log-rank tests were used to visualize differences in progression-free survival between patients within and outside each cluster. Clusters were classified as higher-risk or standard-risk according to the direction of the cluster-specific hazard ratio.

To characterize the biological programs associated with clusters of interest, cluster-specific differentially expressed genes identified using DESeq2 were mapped from Ensembl to Entrez IDs and separated into upregulated and downregulated gene sets. KEGG pathway over-representation analysis was performed independently for each direction using clusterProfiler (v.4.18.4). Disease-specific and unrelated pathway categories were excluded to prioritize molecular and cellular signaling programs. The most strongly enriched pathways were summarized using the enrichment dot plot function from clusterProfiler.

##### 4.3 Identification of cluster-specific noncoding genes associated with outcomes

Cluster-specific differential expression was evaluated using DESeq2 by comparing patients in each transcriptomic cluster with all remaining patients. For candidate ncRNA gene selection, noncoding genes were required to have an adjusted P value below 0.05 and a DESeq2 mean normalized count greater than 10. To prioritize ncRNAs with the strongest cluster-specific expression, candidates were further restricted to genes within the upper 2% of positive $\log_2$fold-change values or the lower 2% of negative $\log_2$fold-change values for each cluster and noncoding genes whose upregulation or downregulation was unique to a single transcriptomic cluster.

The association between each candidate noncoding RNA and progression-free survival was evaluated using a multivariable Cox proportional-hazards model. Standardized variance-stabilized expression was included as a continuous predictor, with adjustment for autologous stem-cell transplantation, race, and sex. Candidates with Cox regression P-values $<0.01$ were retained. Each candidate was provisionally classified as having an oncogene-like or tumor-suppressor-like association by integrating its cluster-specific expression direction, the risk associated with the corresponding patient cluster, and the direction of its expression-associated hazard ratio. For example, cluster-specific ncRNAs upregulated in patient samples from Cluster 9 or Cluster 12 whose increased expression was associated with poorer outcomes were labeled as putative oncogenes, while cluster-specific ncRNAs downregulated in patient samples from Cluster 9 or Cluster 12 whose increased expression was associated with better outcomes were labeled as putative tumor suppressors. Candidate ncRNAs with discordant relationships among these measures were classified as ambiguous and excluded.

##### 4.4 Validating and scoring the final candidate ncRNAs

Candidate noncoding RNAs were evaluated in the integrated single-cell RNA-sequencing atlas to validate their cluster-specific expression in malignant plasma cells. For patients represented in both the bulk and single-cell datasets, plasma-cell expression was averaged by sample to generate sample-level pseudobulk measurements. Expression of each candidate was compared between patients assigned to its associated bulk transcriptomic cluster and all remaining patients using Wilcoxon rank-sum tests with Benjamini-Hochberg correction. Candidates were retained when their cluster-specific differential expression in the single-cell dataset was significant (P-adj. $< 0.05$) and directionally concordant with the bulk RNA-sequencing result. Candidate expression was subsequently compared across major bone marrow compartments, and genes whose highest mean expression occurred in the plasma-cell compartment were retained as the final validated set of 19 ncRNA candidates.

Validated candidates were prioritized using four metrics derived from the bulk RNA-sequencing dataset: cluster-specific $\log_2$fold-change, adjusted differential-expression P-value, magnitude of the progression-free-survival hazard ratio relative to 1, and Cox regression P-value. Each metric was converted to a scaled rank ranging from 0 to 1, with higher values representing stronger evidence. The metrics were combined using weights of 64, 32, 16, and 8, respectively, corresponding to normalized relative weights of 53.3%, 26.7%, 13.3%, and 6.7%. Candidates were ranked in descending order of their weighted composite score.

#### Method S5. *In vitro* quantification of gene expression

##### 5.1 Cell cultures

KMS-12-PE, KMS-18, H929, and MOLP-8 myeloma cell lines were generously provided by Dr. Lawrence Boise's lab (Emory University, Department of Hematology and Medical Oncology). MM cells were cultured in RPMI 1640 media (Gibco, Cat: 11875-093) supplemented with 10% FBS, 100U/mL penicillin/streptomycin (ATCC, Cat: 30-2300), and 1% Glutamax (Gibco, Cat: 35050-061) and were incubated at 37°C and 5% $CO_2$.

##### 5.2 Knockdown and overexpression of *ENSG00000310209*

For knockdown of *ENSG00000310209*, KMS-12-PE, KMS-18, H929, and MOLP-8 cells were transfected with FuGENE HD (Promega, Cat: E231A) and three ENSG00000310209-targeting GapmeRs (Integrated DNA Technologies) labeled antisense oligonucleotide 1 (ASO1), ASO2, and ASO3, and a negative control for each GapmeR treated with FuGENE HD reagent only. The GapmeR sequences can be found in Supplementary Table 1. Knockdown was verified and quantified using RT-qPCR, described below. Knockdown cells were normalized to negative control wells for each GapmeR group. For *ENSG00000310209* overexpression, the ENSG00000310209 sequence was cloned into pCDH-EF1-MCS-BGH-PGK-GFP-T2A-Puro in the Emory Genomic core. Successfully transfected clones were sorted using flow cytometry for GFP to allow for a pure population and were used for downstream analysis.

##### 5.3 Cell proliferation assays

Cell proliferation was assessed using the Dojindo Cell-Counting Kit 8 (Dojindo, Cat: CK04-11). In a 96-well plate, $2.5 \times 10^3$ cells per well were seeded and incubated for 72 hours after transfection with GapmeRs. The proliferation assay was executed following the manufacturer's suggested protocol. Proliferation and viability were assessed using a BMG Labtech CLARIOstar plate reader. Viability assays were conducted in triplicate to ensure scientific rigor and reproducibility.

##### 5.4 Western blot assays

MM cells were lysed using 1X Pierce RIPA Lysis Buffer (ThermoScientific, Cat: 89900) with 1X HaltTM Protease Inhibitor Cocktail, EDTA-free (ThermoScientific, Cat: 78425) for 30 minutes on ice. Protein concentration was determined using PierceTM BCA Protein Assay Kit (ThermoScientific, Cat: 23225) and a BMG Labtech CLARIOstar plate reader. Protein extracts were resolved by SDS-PAGE on Mini-PROTEAN TGX Stain-Free 4-15% 15-well gels (Bio-Rad, Cat: 4568086). Following SDS-PAGE, the gels were transferred onto membranes using Trans-Blot Turbo Transfer Packs (Bio-Rad, Cat: 1704156). Membranes were blocked using EveryBlot Blocking Buffer (Bio-Rad, Cat: 12010020). Membranes were then probed with the following antibodies: β-catenin (Cell Signaling, Cat: 8480T), c-Myc (ABclonal, Cat: A1309), GAPDH (Proteintech, Cat: 60004-1-Ig), and Histone H3 (ABclonal, Cat: A2348). Anti-Rabbit IgG HRP-linked antibody (Cell Signaling, Cat: 7074S) was used for secondary antibody staining. Primary antibodies were diluted in 1X TBST/5% BSA, and secondary antibodies were diluted in 1X TBST/5% dry milk. TBST (BioWorld, Cat: 40120065-2) was used for washing membranes. The signal was developed using SuperSignal West Pico PLUS Chemiluminescent Substrate (ThermoScientific, Cat: 378591), and the signal was detected using a ChemiDoc MP instrument (Bio-Rad, Cat: 12003154).

##### 5.5 RT-qPCR

Total RNA was extracted from MM cell lines using a Quick-DNA/RNATM MiniPrep Plus Kit (Zymo, Cat: D7003) according to the instructions provided by the manufacturer. RNA concentration was assessed using a NanoDrop spectrophotometer. For each experimental group, RT-qPCR was performed using three biological replicates. cDNA was created using the SuperScript VILOTM cDNA Synthesis Kit (Invitrogen, Cat: 11754-050) according to the manufacturer's instructions. The assay was performed using 96-well Hard-Shell PCR plates (Bio-Rad, Cat: HSP9601) with 2X Universal SYBR Green Fast qPCR Mix (ABclonal, Cat: RM21203) and run on a CFX Opus 96 Real-Time PCR System (Bio-Rad, Cat: 12003154). Samples were normalized to negative controls for each experiment, with 18S and β-actin used as housekeeping genes. Fold change was calculated using the Livak method.[25] A list of primers and their sequences can be found in Table S2.
